# Disruption of Mitochondrial Quality Control Genes Promotes Caspase-Resistant Cell Survival Following Apoptotic Stimuli

**DOI:** 10.1101/2021.11.11.468243

**Authors:** Yulia Kushnareva, Vivian Moraes, Julian Suess, Bjoern Peters, Donald D. Newmeyer, Tomomi Kuwana

## Abstract

In cells undergoing cell-intrinsic apoptosis, mitochondrial outer membrane permeabilization (MOMP) typically marks an irreversible step in the cell death process. However, in some cases a subpopulation of the treated cells can exhibit a sublethal response, termed “minority MOMP”. In this phenomenon, the affected cells survive, despite a low level of caspase activation and a subsequent limited activation of the endonuclease CAD (DFFB). Consequently, these cells can experience DNA damage, increasing the probability of oncogenesis. To discover genes affecting MOMP response in individual cells, we conducted an imaging-based phenotypic siRNA screen. We identified multiple candidate genes whose downregulation increased the heterogeneity of MOMP within single cells. Among these were genes related to mitochondrial dynamics and mitophagy, which participate in the mitochondrial quality control (MQC) system. To test the hypothesis that functional MQC is important for reducing the frequency of minority MOMP, we developed an assay to measure the clonogenic survival of caspase-engaged cells. We found that cells deficient in various MQC genes were indeed prone to aberrant post-MOMP survival. Our data highlight the important role of proteins involved in mitochondrial dynamics and mitophagy in preventing apoptotic dysregulation and oncogenesis.

## INTRODUCTION

Apoptosis is a ubiquitous cellular self-elimination process that is critical for the homeostasis of various cell populations. Dysregulated apoptosis is integral to cancer progression and contributes to multiple diseases, including immune and neurodegenerative disorders. Many cancer therapies rely on the enhanced apoptotic death of tumor cells. Apoptosis frequently involves a “cell-intrinsic” pathway involving mitochondria. The central event in mitochondria-dependent apoptosis is mitochondrial outer membrane permeabilization (MOMP), which is induced by BAX and BAK, the key pro-apoptotic BCL-2 family proteins [1, 2]. BAK is constitutively located on the mitochondrial outer membrane (MOM), whereas BAX is mostly soluble in the cytoplasm. When cells receive an apoptotic stress signal, molecules belonging to a subset of the BCL-2 family termed “BH3-only proteins” activate BAX and BAK. Consequently, BAX translocates to the MOM, and both BAX and BAK integrate into the outer membrane and trigger MOMP by inducing the formation of large membrane pores [3-7]. In concert with these BCL-2 family members, additional MOM proteins facilitate BAX-induced pore formation [4]. Anti-apoptotic BCL-2 family members (including BCL-2, BCL-xL, and MCL-1) inhibit MOMP by sequestering the BH3-only proteins and by antagonizing the pore-forming activity of BAX and BAK.

The apoptotic pores in the MOM allow proteins normally residing in the mitochondrial intermembrane space to escape into the cytoplasm. Certain of these proteins are apoptogenic, including cytochrome c, SMAC/DIABLO and OMI/HTRA-1. These proteins induce APAF-1-dependent activation of Caspase-9 and the effector caspases (most prominently Caspase-3), which cleave certain protein substrates that actively promote cell destruction and engulfment [8-10]. Even when caspase activity is blocked, MOMP *per se* usually leads to cell death, as it compromises energy metabolism in the permeabilized mitochondria [11-13]. Although mitochondrial membrane potential and cellular ATP are maintained for some time after MOMP, cells treated with caspase inhibitors typically exhibit a gradual decline in mitochondrial respiration and a complete loss of clonogenic survival over the course of two or three days [12].

Recent reports describe a phenomenon, termed “minority MOMP”, in which cells exposed to relatively weak cell stressors can evade MOMP-dependent death [14-16]. In these atypical cells, only a fraction of the cell’s mitochondria undergoes MOMP. As a result of minority MOMP, small amounts of cytochrome c and SMAC are released into the cytosol, and downstream caspases become activated at sub-lethal levels. This low-level caspase activation leads to a correspondingly limited activation of the apoptotic CAD (DFFB) endonuclease, which enters the cell nucleus and produces a degree of DNA damage. The surviving cells exhibit genome instability and an increased propensity to become oncogenic [17].

The mechanisms controlling the uniformity of MOMP in cells, and hence the frequency of minority MOMP, are unclear. To discover genes that affect MOMP, we conducted a focused high-content siRNA screen that analyzed 1318 genes that had been annotated in databases as having some relationship to mitochondria. A novel aspect of our screen was the identification of genes whose silencing produced a heterogeneous MOMP phenotype in which some mitochondria within the cell were permeabilized, while others remained intact. Our screen yielded functionally diverse candidate genes, including a significant group of genes associated with mitophagy, autophagy, and mitochondrial dynamics, the processes constituting the system of mitochondrial quality control (MQC) [18, 19]. We hypothesized that an increased heterogeneity of MOMP observed in MQC-compromised cells could increase the frequency of minority MOMP [17]. If so, downregulation of MQC-related genes would tend to allow cells to survive stresses that produce a degree of caspase activation, as a result of limited MOMP. To test this hypothesis, we developed a single-cell assay to measure the long-term clonogenic cell survival of cells that had exhibited a measurable degree of caspase activation, in response to treatment with a BH3-mimetic drug. Our results confirmed that deficiencies in various MQC proteins enhanced the survival of caspase-engaged cells. We conclude that mitochondrial dynsmics and mitophagy play a critical role in limiting the frequency of minority MOMP. Thus, proper functioning of the MQC system can be important to prevent oncogenesis that results from an incomplete execution of the mitochondrial apoptosis program.

## MATERIALS AND METHODS

### Cell lines

HeLa cells stably expressing Venus-BAX and OMI-mCherry cells [20] were constructed by the laboratory of Douglas Green (Department of Immunology, St. Jude Children Hospital). U2OS cells with APAF-1 CRISPR knockout (KO) were obtained from Dr. Stephen Tait (Beatson Institute, UK)[21] and Mitofusin (MFN) 1 and 2 double KO (DKO) cells were obtained from Dr. David Chan; California Institute of Technology). Immortalized OPA1 KO and wild type (WT) MEFs were obtained from ATCC (Manassas, VA, USA; deposited by Dr. David Chan). DRP1 KO MEFs were provided by Dr. Stefan Strack (University of Iowa); HeLa cells lacking five mitophagy receptors (TAX1BP1, NDP52, OPTN, NBR1, p62; Penta KO) were provided by Dr. Richard Youle (NIH) [22]. Unless indicated otherwise, cells were maintained in Dulbecco modified Eagles’s medium (DMEM; Life Technologies) containing 10% fetal bovine serum (GeminiBio) and 100 units/ml penicillin/streptomycin at 37°C with 5% CO_2_.

### siRNA screen: cell transfections and treatments

For the primary screen, gene-specific siRNA pools targeting 1318 mitochondria-annotated genes (Table S1) were cherry-picked from a Dharmacon genome-wide siRNA library (siGenome). The list of genes was generated based on a “mitochondria/mitochondrial membranes” queries in the in NIH compiled databases (http://www.ncbi.nlm.nih.gov/gene). Pools of 4 individual gene-specific siRNAs and two non-targeting siRNAs were arrayed in 384-well master plates. Each plate also contained Dharmacon siGenome TOX and siGlo Red oligonucleotides as transfection indicators. Additionally, an siRNA pool targeting BAX was used as a positive control for the inhibition of apoptosis and BAX puncta formation. For reverse transfection, 4.4 µl siRNA picked from each 1 µM stock siRNA solution in the master plate was mixed with 30.6 µl pre-diluted Lipofectamine RNAiMAX transfection reagent (Life Technology). RNAiMAX reagent was diluted 47 times in Gibco Opti-MEM reduced serum medium. The transfection mixtures (35 µl/well) were incubated for 20 minutes at room temperature in a 384-well mixing plate. During the incubation period, Venus-BAX/OMI-mCherry HeLa cells were harvested by trypsinization and resuspended in antibiotic-free DMEM with 10% FBS at 30,000 cells per ml. After incubation, lipid-siRNA complexes were dispensed into triplicate tissue/culture Costar black-wall clear bottom 384-well plates (10 µl/well); siRNAs targeting the same gene were separated in the different plates. Plate handling, transfection reagent/siRNA mixing, and dispensing were performed with Hamilton Star Automated Liquid Handler contained within a Baker Bioprotect class II biosafety cabinet. Cells were added at 40 µl (1200 cells) per well on top of the transfection mixture; final concentration of siRNA was 25 nM in the 50 µl per well volume. The plates were centrifuged at low speed (∼ 300 x *g*) for 1 min and transferred to a humidified CO_2_ incubator (37°C). To minimize plate edge effects, the wells in two rows and columns at the plate edges were not used for transfections and contained only medium with cells. After ∼48 h, the medium was aspirated and replaced with fresh culture medium containing 400 µM etoposide (Sigma-Aldrich) and 20 µM of the caspase inhibitor Q-VD (Q-VD-OPH; SM Biochemicals LLC, Anaheim, CA). Control wells were left untreated (no etoposide). By the time of etoposide treatment, most cells transfected with TOX siRNA had already shown morphological changes consistent with cell death, indicating efficient transfection. Following 24 h of etoposide treatment, the medium was removed, and cells were fixed with 0.5% glutaraldehyde as described previously [23]. After two washes with phosphate-buffered saline (PBS), cells were stained with Hoechst-3342 (Molecular Probes) diluted 1000 times in PBS. Plates were then washed twice with PBS, filled with 50 µl PBS per well, sealed and stored at 4°C. For the secondary screen, 4 individual siGENOME siRNAs per gene were obtained separately from Dharmacon, to target 95 gene candidates chosen from the primary screen. Individual siRNAs were arrayed in replicate 384-well plates, and the experiments were conducted as described above.

### High-throughput image acquisition and analysis

Image collection and processing were done using the Molecular Devices MetaXpress High Content Image Acquisition platform. Images of the cells were acquired in an Image Xpress Micro (IXM) device at 20X magnification with the following filters: DAPI (5060B), for Hoechst nuclear staining; YFP (2427A), for Venus-BAX fluorescence; and TexasRed (4040B), for OMI-mCherry fluorescence. (Numbers indicate the Semrock part number for the filters). Images were collected from 16 sites clustered in the center of the well. In our assay development, the punctate patterns of Venus-BAX translocation and OMI-mCherry retention in mitochondria were analyzed using the granularity module in MetaXpress (version 5.1) software. Individual cells were identified based on nuclear segmentation (DAPI channel images). Z’ factor for the BAX-positive or BAX-negative phenotype assay was calculated as 0.85 based on etoposide-treated and untreated conditions as positive and negative controls, respectively.

High-throughput analysis of primary and secondary screen data was done with a customized algorithm (created as a “journal macro” in MetaXpress) developed for automated phenotype quantification. We configured image analysis to count the percentage of cells that satisfied either of two different criteria: 1) a given cell contains Venus-BAX granules of diameters within the expected diameter range, with total intensity above a certain threshold, or 2) a given cell contains both supra-threshold Venus-BAX and OMI-mCherry granules (not necessarily colocalized) within the appropriate diameter range. The thresholds were adjusted for stringency, to limit false-positive scores from background fluorescence and debris. Plate-to-plate variability was minimal for BAX foci and acceptably low for cells double-positive for BAX and OMI granules.

### Confocal microscopy

Confocal images of Venus-BAX/OMI-mCherry cells were acquired with a 60X oil immersion objective on an Olympus FluoView FV10i automated confocal laser scanning microscope (Olympus Scientific Solutions America Corp, Waltham, MA, USA).

### CRISPR/Cas9-mediated gene depletion

CRISPR experiments were performed using modified synthetic single guide (sg) RNAs from Synthego. Target sequences for guide RNAs were selected with the Synthego CRISPR design tool. RNA oligonucleotides were reconstituted in TE buffer (10 mM Tris-HCl, 1 mM EDTA, pH 8.0) according to recommendations from the manufacturer. Ribonucleoprotein (RNP) complexes were formed from sgRNA and recombinant Cas9 2NLS protein (New England Biolabs) mixed at a sgRNA to Cas9 ratio of 4.5: 1. After a 20-30 min incubation at room temperature, assembled RNP were delivered into cells by electroporation using a ThermoNeon™ device and 10 µM tips (ThermoFisher). For one sample (∼2×10^5^ cells), 3 µl of sgRNA (from 30 mM stock solution) were mixed with 1.5 µl Cas9 protein (from 20 µM stock solution). U2OS cells were electroporated at 1230 v/10ms/4 (pulse voltage, width, pulse number) settings. Immediately after electroporation, cells were transferred to 12-well plate with pre-warmed antibiotic-free DMEM media with 10% FBS. After two-three days of incubation, a portion of the control (unedited) and CRISPR cells were harvested for genomic DNA isolation using a QIAGEN genomic DNA purification kit. The remaining cells were left in culture for further propagation and clonal selection. Genomic DNA concentrations were measured using a NanoDrop instrument (ThermoFisher). Primer design and PCR amplification of the edited region were done according to Synthego recommendations. Sanger sequencing of PCR amplicons was performed at Genewiz or EtonBio; the resulting DNA sequencing chromatograms were analyzed using the Inference of CRISPR Edits (ICE) algorithm (Synthego). ICE analysis of CRISPR-edited genomic regions typically demonstrated at least 70% editing efficiency (i.e. 70% knockout cells in the pool); sgRNA sequences used are shown **Fig. S2**. For clone selection, CRISPR cells were expanded for 2-3 additional days, harvested and subjected to sorting into 96-well plates using BD FACSAria-3 or FACSAria-4 Fusion instruments; typically, 1-4 single cells were dispensed into one well. After clonal expansion, cells were analyzed for knockout efficiency as described above. For each gene of interest, two clones with verified knockout were combined for use in further experiments.

### Assay for clonogenic survival after caspase activation via MOMP (outlined in Fig. 2)

Cells (CRISPR knockout U2OS cells and MEFs (knockout and matched WT) were plated out at 3×10^5^ per well containing 1 ml media in 6-well plates the day before the experiment, 2 wells per condition. The cells were then treated with ABT-199 or ABT-737 at 2.5, 5 or 10 µM for 5 h. In the last 30 min of incubation, a caspase reporter dye, CellEvent Caspase3/7 Green (Life Technology; R37111), was added at 30 µl per well. Cells were harvested with Trypsin/EDTA and washed with medium and PBS. Finally, cells were resuspended in 300 µl of sorting buffer consisting of 1% BSA in PBS containing 20 µM of Q-VD, where Q-VD was included to stop the caspase reaction. We noticed that, when these caspase-activated cells were left at room temperature or 4°C for more than 30-40 min, their ability to survive was compromised. Therefore, care was taken to sort the cells immediately after harvesting. To minimize artifacts from sample handling delay, we operated two cell sorters (FACSAria; BD) simultaneously for wild type and gene-defective cells. Also, the sorting chamber and the plate holder were kept at 37°C during the run to prevent temperature shock to the cells. Six hundred cells within the population gated for green fluorescence were sorted in duplicate into a 48-well plate containing 300 µl of conditioned medium per well. The cells were grown for 8-12 days, during which an additional 500 µl of conditioned medium was added at day 4-5. Colonies were then stained with 6% glutaraldehyde containing 0.5% crystal violet. Non-treated cells distributed in the non-green gate were sorted as above and used as a control representing 100% growth. Because individual cell colonies tended to merge over time, we measured the total area occupied by colonies, using the ImageJ macros developed by Guzman et al [24]. The percentage of surviving caspase-engaged cells in the total population was normalized against the 100% growth control. The data are summarized from 2 to 3 independent experiments, as indicated. P values were calculated using the mean and the standard deviation in each set of KO and WT cells with two-way ANOVA analysis using Prism 7 (Graphpad Software).

## RESULTS AND DISCUSSION

### Screening strategy and assay development

To identify novel cell-intrinsic regulators of MOMP, we designed a high-content image-based siRNA screen. As we were primarily interested in mitochondrial function, we carried out a focused siRNA screen targeting genes with annotated relationships to mitochondria (**Table S1**). Our strategy took advantage of a HeLa cell line expressing two fluorescent reporter proteins (described previously by Llambi et al [20]) that enabled us to simultaneously interrogate both BAX activation and MOMP. In living cells, Venus-BAX is cytoplasmic (diffuse), and OMI-mCherry is localized to mitochondria. When apoptosis is induced, Venus-Bax is translocated to mitochondria, and OMI-mCherry is released into the cytoplasm and degraded. Normally, once apoptotic BAX translocation is initiated in a given cell, mitochondrial intermembrane space proteins (including cytochrome c or OMI) are released in a synchronous, rapid fashion [14, 20, 25]. The release of proteins from the mitochondrial interior can only be restricted under special circumstances, e.g. cristae junction remodeling by overexpression of a mutant form of the optic atrophy 1 (OPA1) protein [26] or MOMP inhibition by a recently described endolysosome-linked mechanism [27]. However, in some cells the release of intermembrane proteins is not an all-or-none event and mitochondria within the same cell display heterogeneity in MOMP response [16, 17]. We hypothesized that downregulation of certain genes could increase MOMP heterogeneity and promote post-MOMP cell survival, even in caspase proficient cells.

To begin to test our hypothesis, we analyzed punctate mitochondrial distribution of Venus-BAX and OMI-mCherry in cells treated with the apoptosis inducer, etoposide. The caspase inhibitor Q-VD was included in these experiments to decrease premature cell loss and detachment. We configured our automated image analysis to quantify three cellular phenotypes: 1) apoptotic cells with mitochondria that have undergone MOMP – containing translocated BAX and lacking intramitochondrial OMI (BAX puncta-positive); 2) non-apoptotic cells with intact mitochondria lacking BAX and containing OMI (BAX puncta-negative), and 3) atypical cells with mitochondria containing BAX but which have not released OMI (BAX/OMI puncta double-positive cells) (**Fig 1. A, B**). Untreated (no etoposide) cells transfected with control siRNA were non-apoptotic and showed normal mitochondrial morphology (**Fig. 1A, E**) indicating that our optimized transfection conditions produced minimal toxicity. As expected, most etoposide-treated cells fell into either of the first two categories, while BAX/OMI double-positive cells were minimally present (**Fig.1 A, E**). In assay validation experiments, we tested the effect of OPA1 siRNA. OPA1 mediates mitochondrial fusion in conjunction with its cleavage by the mitochondrial protease OMA1 [28-30] and has multiple roles in apoptotic signaling [23, 26, 31-33]. Knockdown of OPA1 protein increased the number of BAX/OMI double-positive cells compared with cells transfected with a non-targeting siRNA (**Fig. 1F)**. Furthermore, high-resolution confocal imaging of OMA1 siRNA-treated cells revealed mitochondrial heterogeneity in the MOMP response: in individual BAX/OMI double-positive cells, some mitochondria contained translocated BAX (green puncta) and lost OMI, whereas other mitochondria lacked BAX and retained OMI (red puncta) (**Fig. 1D**).

**Figure 1.**
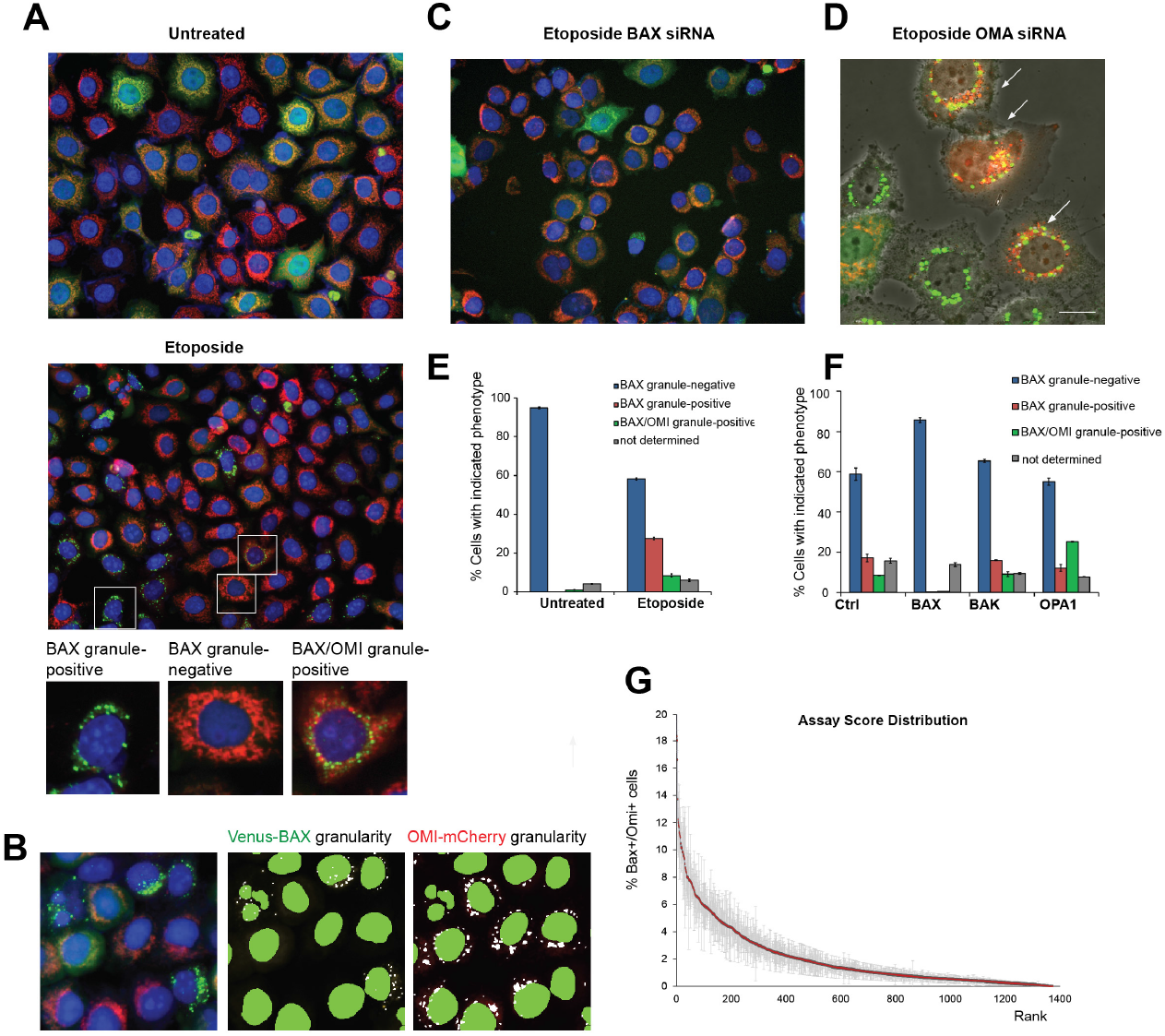
Imaging-based screening assay for regulators of apoptotic MOMP response. (**A**) Representative epifluorescence microscopy of untreated and etoposide-treated cells expressing Venus-BAX (green; YFP channel) and OMI-mCherry (red; Texas Red channel). Bottom panel illustrates enlarged cells with indicated phenotypes. Nuclei were stained with Hoechst 33342 (blue; DAPI channel). (**B**) Identification of Venus-BAX and OMI-mCherry puncta using MetaXpress granularity application module. A representative enlarged image of etoposide-treated cells (left panel) and corresponding image segmentations. The granularity module identifies Venus-BAX (middle panel) and OMI-mCherry puncta. Note that cells with BAX “granules” do not contain OMI “granules”, and *vice versa*. Nuclear segmentation settings correctly identify fragmented (apoptotic) and normal size nuclei. (**C**) BAX siRNA inhibits Venus-BAX puncta formation and OMI-mCherry release in etoposide-treated cells. (**D**) A confocal image of live Venus-BAX/OMI-mCherry cells transfected OMA1 siRNA. Arrows indicate cells with heterogeneous MOMP. (**E, F**) Examples of phenotype quantification using granularity application module and high-throughput microscopy. (**E**) Quantification of indicated phenotypes in untreated and etoposide-treated cells transfected with a non-targeting (control) siRNA; “non-determined” phenotype corresponds to a small fraction of non-fluorescent cells. (**F**). Effects of indicated siRNAs on the phenotype distribution. Data are mean ± S.E.M. (n=3 replicate wells). (**G**) Sorted assay scores for the primary screen siRNA set in triplicate plates. Numbers indicate the percentage of cells positive for both Venus-BAX and OMI-mCherry foci. Values shown are mean and SEM, n = 3. Hits were identified from the top 10% tail of the BAX/OMI score.

A BAX siRNA pool (which targets both endogenous BAX and ectopic Venus-BAX expression) potently inhibited etoposide-induced apoptosis and markedly reduced Venus-BAX fluorescence (**Fig. 1C, F)**. However, knockdown of BAK did not inhibit apoptosis, indicating that in these etoposide-treated cells, MOMP was predominantly BAX-dependent. Transfection efficiency in high-throughput screening experiments was verified using a cell death-inducing transfection marker (siTOX) that consistently produced ∼90% loss of cells within 2 days post-transfection (not shown). Efficient knockdown of other proteins produced by siRNA pools was confirmed in supplementary experiments (**Fig. S1A**).

### The siRNA screen uncovers potential regulators of MOMP

The image-based assay was used to screen 1318 gene-specific siRNA pools. We ranked gene scores in three groups corresponding to the cell phenotype categories noted above: genes whose knockdown I) increased the percentage of Venus-BAX puncta; II) decreased the percentage of cells with Venus-BAX puncta; or III) increase the percentage of BAX/OMI double-positive cells. When we applied a 10% cut-off in the score distribution range, we identified ∼200 initial “hits” (with scores ∼3-5 fold above or below controls for the Categories I and II). Hits in the Category III were from the top 10% tail of the score distribution (**Fig. 1G**). For a secondary screen, we chose to re-assay 95 of the primary screen hits, based on their effect scores and our judgment concerning their biological interest (we decided not to pursue some “housekeeping” genes.) In this secondary screen, we tested the four siRNAs from each siRNA pool individually. To reduce the likelihood of off-target effects, we required at least 3 of the 4 individual siRNAs to give concordant results [34]. Applying this more stringent criterion yielded a final list of 63 candidate genes (**Table S3**). Several hits in Categories I and II were consistent with previous reports (**Table S2**). For example, the effect of siRNA targeting BNIP3 (Category II) is consistent with the known cell death-promoting activity of this protein [35]. Acting in the opposite manner, the siRNAs scoring in Category I increased the percentage of cells undergoing MOMP. For example, several candidate genes in this group are required for metabolism of the mitochondrial lipid, cardiolipin (PLSCR3, PRELID1, and TRIAP1). Although cardiolipin is important for BAX pore formation [6, 36], in certain paradigms cardiolipin deficiency potentiates the release of apoptogenic proteins [37, 38]. In particular, the deficiency of p53-regulated protein TRIAP1 impaired cardiolipin level in mitochondria, compromised bioenergetics and potentiated cytochrome c release [38]. Potential novel regulators of MOMP include metabolic enzymes ACADL (long-chain acyl-coenzyme A dehydrogenase) which catalyzes one of the early steps in the circle of mitochondrial beta-oxidation of fatty acids, and FAHD1 (fumarylacetoacetate hydrolase domain-containing protein) with putative oxaloacetate decarboxylase activity in mitochondria. The roles of these proteins in mitochondrial metabolism and cell senescence are emerging [39, 40] and their inhibitory effects on MOMP could be of interest for further investigations. In this study, we did not further pursue the hits in the Categories I and II but focused on the unique phenotype produced by the hits in category III. Of note, in nearly all cells containing both BAX and OMI puncta, BAX and OMI were not colocalized, implying that some mitochondria within a cell had undergone MOMP, while others had not (similar to the phenotype shown in **Fig. 1D**). Perhaps the most striking outcome of our screen is that a majority of hits in the Category III are related to the mitochondrial quality control (MQC) system.

### The interplay of MQC and MOMP

The concept of MQC is that mitochondrial dynamics (fission and fusion) work in tandem with mitophagy (the autophagic elimination of dysfunctional mitochondrial fragments) to maintain mitochondrial structural and functional integrity. If defective mitochondria cannot be repaired through fusion with functional organelles, they are prone to excessive fragmentation and elimination by mitophagy. It is postulated that asymmetrical fission segregates defective mitochondria, making the mitochondrial population inherently heterogeneous. These isolated small mitochondria are then redirected into a pre-autophagic pool [41, 42]. In cells dysfunctional for mitochondrial dynamics or mitophagy, damaged mitochondria would be predicted to accumulate, and the organelles would become abnormally heterogeneous with respect to bioenergetic function and the distribution of proteins involved in MOMP [43, 44]. Our screen tended to confirm this prediction.

### Downregulation of mitophagy-related genes promotes MOMP heterogeneity

Among the hits in Category III, both ubiquitin-dependent (e.g. ATG12, MUL1) and ubiquitin-independent (e.g. FUNDC1, BNIP3L, BLOC1S1/GCN5L1, TRIAP1, MARCH5) mitophagy mechanisms [45-48] were represented (Table 1). ATG12 is ubiquitin-like protein that is integral for general autophagy and has additional roles in apoptosis and cell survival [49]. For example, ATG12 deficiency compromises mitochondrial function and promotes cellular oncogenic transformation [50]. One of the top-scoring hits was BLOC1S1/GCN5L1, which is reported to have multiple functions, including the coordinate regulation of mitochondrial biogenesis and mitophagy, through protein acetylation [51]. BNIP3L/NIX (originally described as a BCL-2-related pro-apoptotic protein) and FUNDC1 directly interact with the LC3 protein in the autophagosome formation pathway and promote mitophagy facilitated by hypoxic conditions [48, 50, 52, 53].

**Table 1.** A list of genes whose downregulation increased mitochondrial heterogeneity (BAX/OMI-positive phenotype) in siRNA screen. MQC functions (mitochondrial dynamics, mitophagy) of the hits are highlighted in red. Other highlighted functions that can affect mitochondrial heterogeneity include regulation of mitochondrial respiration and apoptosis. Genes in boldface (RMDN3, ATG12, and BNIP3L) were selected for further experiments. Numbers indicate assay scores obtained in the secondary screen. Other secondary screen hits are listed in Table S2.

| Gene | score | Function |
| --- | --- | --- |
| RNF5 | 85.3 | E3 ubiquitin ligase; ER quality control; translocates from ER to mitochondrial in antiviral response; controls UPR, **autophagy**, anti-tumor immunity [69-72] |
| LETM1 | 78.8 | Mitochondrial Ca<sup>2+</sup> and/or K<sup>+</sup> transport; implicated in assembly of ETC supercomplexes and maintenance of tubular **mitochondrial morphology** [64, 73, 74] |
| MARCH5 | 64 | E3 ubiquitin ligase, regulates **mitophagy** and **mitochondrial dynamics** via targeting FUNDC1, MFN1/2 and Fis1; involved in MAVS signaling [75-77] |
| FUNDC1 | 59.3 | Regulation of LC3-dependent **autophagy** and **parkin-independent mitophagy**; a MARCH 5 target; regulates **MQC** in hypoxia, **interacts with Drp1** [46, 75, 78] |
| DISC1 | 50.3 | Disrupted-in-schizophrenia-1; enriched in MAM; controls Mitofilin stability, regulates **mitochondrial trafficking**, Ca<sup>2+</sup> dynamics, and **bioenergetics** [79-81] |
| ULK1 | 48.3 | **Autophagy** activating kinase, phosphorylates FUNDC1; interacts with Bcl-2-L13 and LC3B to induce **mitophagy**; negatively regulated by mTOR [82-84] |
| ATG12 | 42.8 | Integral component of general and selective **autophagy**, binding partner of ATG5; promotes **apoptosis**; regulates **mitochondrial biogenesis** [14, 85, 86] |
| RMDN3 | 39.5 | Encodes tyrosine phosphatase-interacting protein-51 (PTPIP51); controls mitochondria-ER tethering, ER stress, **autophagy**, **parkin-dependent mitophagy** [54-56] |
| NR2C2 | 38.7 | Nuclear receptor/transcriptional regulator (also known as TR4); DNA repair function; KO promotes tumorigenesis, **complex I deficiency**, myopathy [87-89] |
| IER3 | 37 | Early response gene, multifunctional; positive and negative **apoptosis** regulator, tumor-suppressive activity; interacts with Mcl-1; **modulates complex V** [90-92] |
| SLC25A14 | 33.3 | Encodes UCP5; mitochondrial metabolite/anion transporter; KD **decreases $\Delta\Psi$**; implicated in ROS production, and DJ-1-dependent neuroprotection[93-95] |
| PHB | 33.3 | Prohibitin, controls **ETC protein stability**, **OMA1/OPA1-dependent fusion** and cristae morphogenesis, **apoptosis**, cancer cell proliferation [96-98] |
| MRPL12 | 33 | Mitochondrial ribosomal protein, regulates mitochondrial gene expression; upregulated in cancer; mutations are linked to a mitochondrial disease [99-101] |
| BNIP3L | 30 | Atypical **pro-apoptotic** Bcl-2 family protein, multifunctional, mediates hypoxia-activated **autophagy and mitophagy** with effects on **mitochondrial fission** [48, 52, 102] |
| GPER1 | 24 | G-protein coupled estrogen receptor |
| TMEM127 | 24 | Endosome-associated tumor suppressor gene, linked to various cancers; regulates **Rab5-mediated endosomal fusion** [103]; mitochondrial function is unknown |
| ATCAY | 23.7 | Encodes caytaxin, a brain-specific kinesin interacting protein; involved in kinesin-dependent **mitochondrial motility** on microtubules [104] |
| TRIAP1 | 22 | p53-regulated **antiapoptotic factor**, linked to tumorigenesis; KD promotes **MOMP and fission**; lipid transfer activity, cardiolipin biosynthesis [38, 105, 106] |
| C19orf12 | 16.3 | an orphan mitochondrial protein whose loss of function is linked to neurodegenerative diseases; involved in lipid metabolism, stimulates **autophagy** [107, 108] |
| SPATA19 | 15 | Spermatogenesis associated 19; important for sperm motility by regulating mitochondrial structure and function [109] |
| BLOC1S1 | 9.5 | General control of amino acid synthesis 5 like-1 (GCN5L1/BLOC1S1); endosome-lysosomal function; mitochondrial protein acetylation and **mitophagy** [51] |
| TRAC1 | 7 | Essential for kinesin 1-depencent **mitochondrial trafficking** [57, 58] |

### Other MQC-related hits in the Category III

Besides well-defined components of mitophagy pathways, our screen yielded multiple candidates that could affect MQC as part of their function. For example, PTPIP51 (encoded by the RMDN3 gene) was shown to be important for mitochondria-endoplasmic reticulum (ER) tethering, and therefore, its putative role in MQC and apoptosis could be linked to multiple functions regulated by ER-mitochondrial contacts, such as ER stress-induced Ca^2+^ release [54], autophagosome formation and mitochondrial fission [55, 56].

Another functional group represented by multiple hits in the screen (ATCAY, DISC1, TRAK1 and SPATA19) involves mitochondrial motility on microtubules, in which motor proteins such as Kinesin 1 interact with certain mitochondrial membrane proteins, e.g. MIRO1 [57, 58]. We suspect that these genes appeared as hits in our screen because microtubule-based motility is important for mitochondrial fusion, including the transient fusion event known as “kiss and run” [59, 60]. Since fusion is a key element of MQC, a loss of mitochondrial motility would be expected to increase the functional heterogeneity of the mitochondrial population within each cell.

Other interesting candidates include multifunctional E3 ubiquitin ligases RNF5 and MARCH5. Mitochondrial targets and pathways regulated by RNF5 remain to be identified. MARCH5 reportedly regulates activities of mitofusin-1 (MFN1) and dynamin-related protein 1 (DRP1) or its outer membrane receptor MFF, the key mediators of mitochondrial fusion and fission, respectively [61-63]. However, DRP1 and MFN themselves did not show significant effects in our screen, perhaps due to the redundancy. Among multifunctional proteins that may influence mitochondrial dynamics indirectly, LETM1 is the mitochondrial Ca^2+^/H^+^ antiporter with pleiotropic effects on ATP production, mitophagy, cristae structure and mitochondria morphology [64]. In particular, mitochondria lacking LETM1 are prone to undergo DRP1-independent fission [65]. In our screen, LETM1 knockdown also increased mitochondrial fragmentation (not shown) as well as MOMP heterogeneity. Other candidate genes in the Category III are highlighted in Table 1. Overall, our screen suggests that a deficiency in MQC disturbs the normal apoptotic response and, in particular, promotes heterogeneous MOMP.

### An assay to measure clonogenic cell survival despite caspase activation

Based on the number of hits our screen being involved in MQC, we hypothesized that MQC-dependent mitochondrial heterogeneity could result in an increased frequency of “minority MOMP”. In this scenario, when a minor fraction of mitochondria undergoes MOMP, the resulting sub-lethal caspase activation leads to DNA damage and promotes oncogenesis [17]. To test our hypothesis, we developed a clonogenic survival assay based on that described by Ichim et al. [17] (**Fig. 2**). In this assay, we induced apoptosis with a BH3-mimetic compound, ABT-737 or ABT-199, incubated the cells with a fluorescent caspase-reporter compound, and used FACS to sort out cells that exhibited caspase activation. We then plated the cells and, after several days in culture, quantified the percentage of cells that survived and formed colonies.

**Figure 2.**
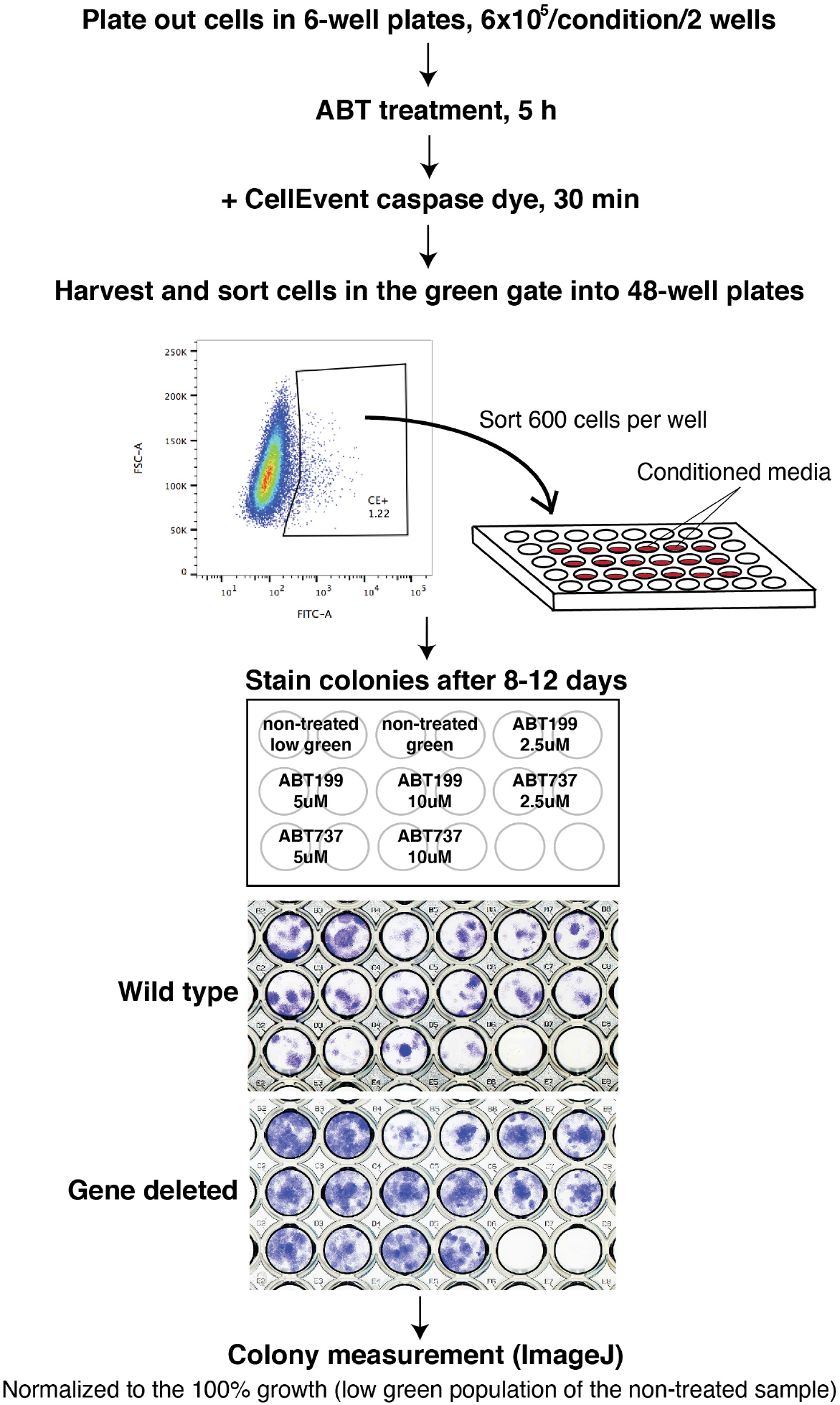
Cell-based ‘minority MOMP’ assay. Cells were treated with ABT199 or ABT737 for 5 h and a caspase dye was added for the final 30 min. Cells were harvested and 600 cells from the FITC-positive gate were sorted into 48-well plates containing the conditioned media. Grown colonies were stained with crystal violet and colony areas were measured using the macro developed by Guzman et al. [24](see Methods)

If the caspase dye faithfully reports MOMP-induced caspase activation, rather than unspecific staining and/or caspase activation independent of MOMP, we would predict that cells deficient in caspase activation via the intrinsic pathways would not display cell survival in this assay. To confirm this, we compared APAF1 CRISPR KO U2OS cells to their parental WT control cells. APAF1 is required to activate Caspase-9 and subsequently the effector Caspases-3 and −7, following the MOMP-dependent release of cytochrome c, SMAC and OMI. As shown in **Fig 3**, only a trace amount of cells were stained by the dye in the APAF1 KO cell population treated with a BH3 mimetic, ABT-737 (**Fig. 3A**). In contrast, the WT cells showed a population of cells stained with the dye. The same number of dye-stained cells in the gate was collected and subject to a colony formation assay. The percentage of the surviving cells in the entire population was then calculated. APAF1-deficient cells from this gate did not survive clonogenically (**Fig. 3B**), suggesting that the staining in these cells is not a result of genuine caspase activation. We conclude that our clonogenic assay reflects true MOMP-dependent survival of caspase-engaged cells and thus indicates the occurrence of minority MOMP.

**Figure 3.**
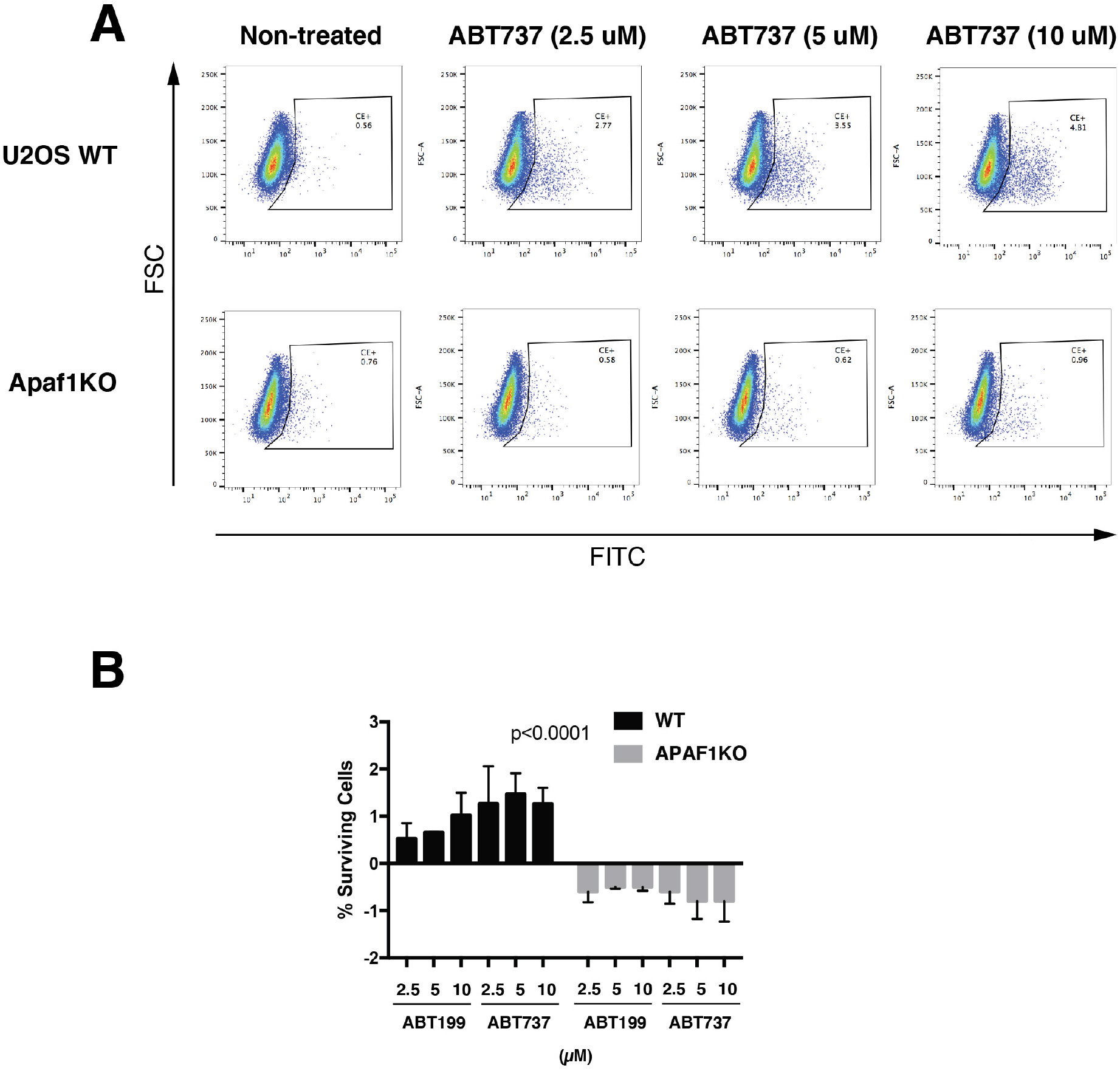
Survived cells had APAF1-dependent caspase activation. **(A)** Dot-plots of wild-type U2OS and APAF1 CRISPR KO cells stained with a caspase dye (CellEvent; Life Technologies) after treatment with ABT737 at 2.5, 5 or 10 µM for 5 h. (**B**) Calculated percentages of surviving cells in WT U2OS cells and APAF1 KO are shown. Data presented are from two independent experiments.

### MQC deficiency promotes minority MOMP

To test further whether MQC is important to mitigate minority MOMP, we used the clonogenic assay described above to investigate the effects of deleting selected candidate genes from our screen as well as other known MQC factors. We generated CRISPR cells (U2OS) with perturbed expression of ATG12, RMDN3 (PTPIP51) and BNIP3L. Knockdown of each of these genes produced MOMP heterogeneity in our imaging assay (**Table 1**). To obtain stable knockout cells, we generated CRISPR-edited cell pools and derived clonal populations from single cells (as described in Methods). Validation of the CRISPR-based gene editing efficiency is shown in **Fig. S2**. As shown in **Fig. 4**, we found that cells depleted of either RMDN3 or ATG12 showed significantly higher post-MOMP survival than WT cells. However, BNIP3L KO did not show an increase in minority MOMP. Reasons for this are unknown but could be related to other apoptosis-related activities of BNIP3L.

**Figure 4.**
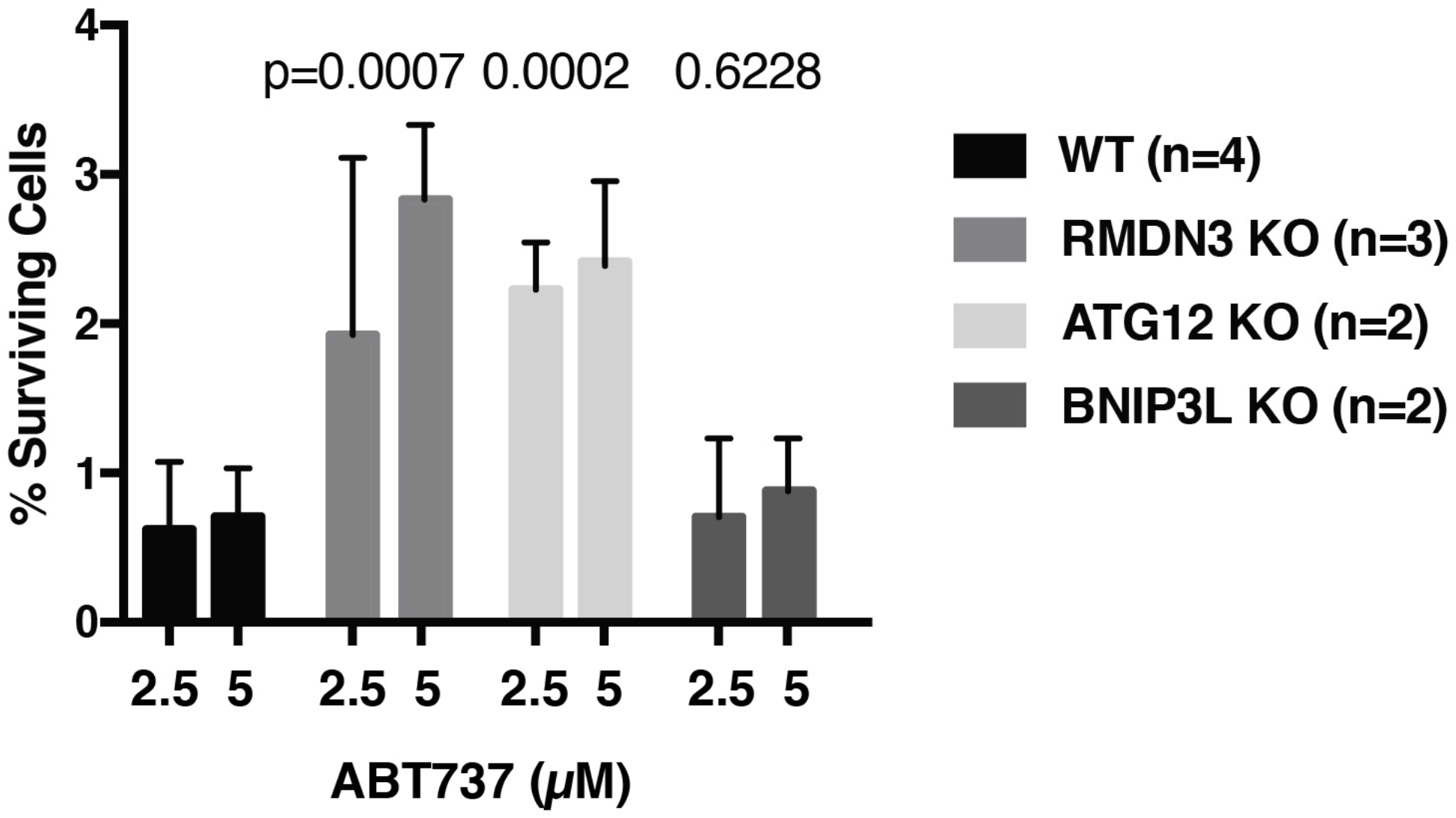
Heterogeneous MOMP induced more survival of cells after caspase activation. Candidate genes from the screen for the BAX/OMI-positive phenotype (RMDN3, ATG12 and BNIP3L) were chosen and deleted by CRISPR in U2OS cells and subjected to the clonogenic survival assay. The percentages of surviving cells were plotted. P values above the bar graphs show the difference in survival between the WT U2OS cells and each KO line.

To examine further the hypothesis that compromised MQC promotes minority MOMP, we performed experiments using previously derived cells in which genes known to be directly involved in mitochondrial dynamics or mitophagy had been deleted, namely MFN1 and 2 double knockout (DKO), OPA1 KO, DRP1 KO MEFs and “penta-KO” HeLa cells that had been shown to be deficient in mitophagy due to the deletion of 5 mitophagy receptor genes [22]. As shown in **Fig. 5**, all of these MQC-deficient cells exhibited an enhancement of cell survival in our clonogenic assay, compared with their matched controls. (The effect in DRP1 KO cells was not statistically significant; in this case, a weaker effect could reflect the slowing of cell proliferation that was reported with DRP1 deletion [66]. Thus, the loss of these genes important for MQC promoted the proliferative survival of caspase-engaged cells, again implying an increased frequency of minority MOMP.

**Figure 5.**
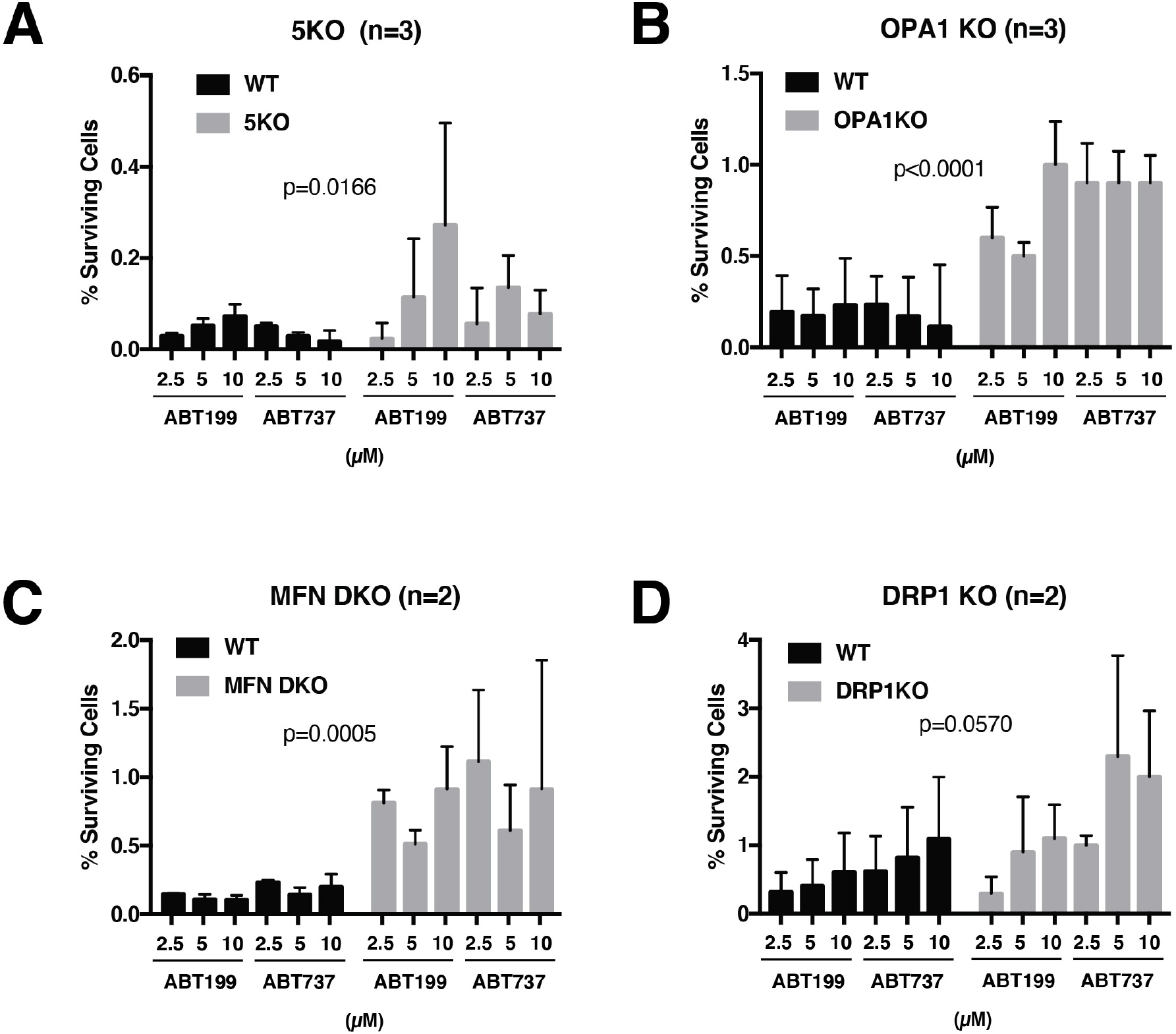
Minority MOMP is enhanced by deletion of mitophagy- or mitochondria dynamics-related genes. (A) Mitophagy defective penta-KO cells and their parental WT HeLa cells. (**B**) OPA1 KO MEFs and their matched WT MEFs. (**C**) MFN DKO MEFs and their matched WT MEFs. (**D**) DRP1 KO MEFs and DRP1-positive WT MEFs used in the MFN set in **C**. n denotes the number of independent experiments that were averaged. Error bars represent standard deviation.

## Conclusion

Taken together, our data allow us to conclude that the MQC system is critical for mitigating the phenomenon of minority MOMP. This result has implications for oncogenesis: even a small increase in cell viability resulting from sublethal caspase activation could potentially raise the frequency of oncogenic cell transformation [67, 68]. Therefore, treatments developed to limit minority MOMP could improve the cytotoxic effects of anti-cancer treatments. For example, proteins that are known to inhibit mitophagy, such as SIAH3 [22], are potential therapeutic targets. Also, we predict, based on our observation that placing sorted caspase-engaged cells on ice eliminated their ability to survive clonogenically (see Methods), that cryotherapy could prevent minority MOMP when used in combination with pro-apoptotic cancer therapeutics such as BH3 mimetic drugs. In conclusion, our study provides new insights into the mechanisms of aberrant apoptosis that are important for developing therapeutic strategies.

## ACKNOWLEDGMENTS

We thank the Flow Cytometry Core Facility at La Jolla Institute for their expert help in developing the clonogenic assay. We thank Drs. Stephen Tait (Cancer Research UK Beatson Institute, University of Glasgow, UK), Sonia Sharma (Functional Genomics Center at LJI) and Alexander Andreyev (The Scripps Research Institute) for helpful discussions and critical reading of the manuscript. This work was supported by R21CA216304 and a SPARK award from the LJI Board of Directors to D.D.N.

