## Supplemental Table S1, S2, Figure S1, S2 for "Disruption of Mitochondrial Quality Control Genes Promotes Caspase-Resistant Cell Survival Following Apoptotic Stimuli"

**Table S1**. **Mitochondria-annotated genes targeted by siGENOME siRNA pools in the primary screen**. The list indicates gene names, Entrez gene ID and corresponding siRNA catalog numbers.

**Table S2**. **Secondary screen results summary**. Numbers show assay scores (% of cells) for top candidates in each of the three phenotype categories. Genes in boldface (RMDN3, ATG12, and BNIP3L) were selected for further experiments.

**Figure S1. Knockdown efficiencies of selected siRNA pools in Venus-BAX/OMI-mCherry HeLa cells**. Cells were reverse- transfected with a non-targeting (nt) and indicated siRNAs at 25-50 nM. Transfection conditions were essentially as described in Methods for screening experiments but scaled up to 6-well plate format to obtain sufficient amounts of cells for western blot analysis. TOM20 was used as loading control.

**Figure S2. CRISPR/Cas9-mediated knockout of selected gene candidates. (A)** Single guide RNA sequences and CRISPR-mediated gene editing efficiencies in U2OS cells. (**B**) An example of ICE analysis of CRISPR-edited genomic regions shown for RMDN3 knockout cell pool and clone. Other details of the experiments are described in Methods.

### Table S1

| | | | | | | | | |
|---|---|---|---|---|---|---|---|---|
| AARS | 16 | M-011565-01 | ANXA7 | 310 | M-010760-01 | BCL2L2 | 599 | M-004384-02 |
| AASS | 10157 | M-009247-00 | APBB1 | 322 | M-011213-03 | BCR | 613 | M-003875-05 |
| ABCA12 | 26154 | M-008407-01 | APC | 324 | M-003869-01 | BDH | 622 | M-008643-02 |
| ABCA4 | 24 | M-009533-02 | APEX1 | 328 | M-010237-01 | BECN1 | 8678 | M-010552-01 |
| ABCA8 | 10351 | M-008347-00 | APEX2 | 27301 | M-013730-00 | BEST1 | 7439 | M-019825-01 |
| ABCB10 | 23456 | M-007300-00 | APOE | 348 | M-006470-00 | BID | 637 | M-004387-02 |
| ABCB11 | 8647 | M-007301-00 | APOO | 79135 | M-014318-00 | BIK | 638 | M-004388-02 |
| ABCB4 | 5244 | M-007302-01 | APOOL | 139322 | M-027214-01 | BIRC2 | 329 | M-004390-02 |
| ABCB5 | 340273 | M-007303-01 | AREG | 374 | M-017435-00 | BIRC3 | 330 | M-004099-02 |
| ABCB6 | 10058 | M-007304-00 | ARL13B | 200894 | M-017365-00 | BIRC4 | 331 | M-004098-01 |
| ABCB7 | 22 | M-007305-00 | ARL2 | 402 | M-011585-01 | BLOC1S1 | 2647 | M-012580-00 |
| ABCB8 | 11194 | M-007306-01 | ARL2BP | 23568 | M-013074-00 | BLOC1S2 | 282991 | M-018342-00 |
| ABCC9 | 10060 | M-007316-01 | ARL6 | 84100 | M-017298-00 | BMF | 90427 | M-004393-04 |
| ABCD1 | 215 | M-009605-01 | ARL6IP5 | 10550 | M-012229-00 | BNIP1 | 662 | M-011222-01 |
| ABCD3 | 5825 | M-009909-01 | ARMCX3 | 51566 | M-013326-00 | BNIP3 | 664 | M-004636-01 |
| ABCE1 | 6059 | M-008702-01 | ASAH2 | 56624 | M-005229-00 | BNIP3L | 665 | M-011815-01 |
| ABCF2 | 10061 | M-009286-00 | ASS | 445 | M-010257-03 | BOC | 91653 | M-008413-00 |
| ABCG2 | 9429 | M-009924-01 | ATAD3A | 55210 | M-008191-01 | BOK | 666 | M-004394-00 |
| ABL1 | 25 | M-003100-02 | ATAD3B | 83858 | M-019443-01 | BRI3BP | 140707 | M-017153-00 |
| ACAA2 | 10449 | M-008773-00 | ATCAY | 85300 | M-027139-01 | BRP44 | 25874 | M-020233-01 |
| ACACB | 32 | M-004759-02 | ATF2 | 1386 | M-009871-00 | BRP44L | 51660 | M-020459-00 |
| ACAD9 | 28976 | M-009002-00 | ATG12 | 9140 | M-010212-02 | C12orf62 | 84987 | M-015041-00 |
| ACADL | 33 | M-009851-00 | ATG4D | 84971 | M-005790-01 | C14ORF112 | 51241 | M-021166-01 |
| ACADS | 35 | M-010041-02 | ATG5 | 9474 | M-004374-04 | C14ORF160 | 79944 | M-008130-01 |
| ACADVL | 37 | M-009392-02 | ATP1A1 | 476 | M-006111-02 | C14ORF2 | 9556 | M-019882-01 |
| ACAT1 | 38 | M-009408-00 | ATP1B1 | 481 | M-008381-00 | C14ORF68 | 283600 | M-007317-01 |
| ACIN1 | 22985 | M-014157-01 | ATP2A1 | 487 | M-006113-00 | C17ORF35 | 8834 | M-005440-01 |
| ACSL1 | 2180 | M-011654-00 | ATP2B2 | 491 | M-006116-00 | C18orf55 | 29090 | M-020711-01 |
| ACSL3 | 2181 | M-010061-00 | ATP5A1 | 498 | M-017064-01 | C19ORF12 | 83636 | M-014731-01 |
| ACSL4 | 2182 | M-009364-00 | ATP5B | 506 | M-018615-01 | C1orf151 | 440574 | M-033619-00 |
| ACSL5 | 51703 | M-006327-00 | ATP5C1 | 509 | M-012705-01 | C1orf166 | 79594 | M-007062-02 |
| ACSL6 | 23305 | M-007748-01 | ATP5D | 513 | M-017852-01 | C1orf31 | 388753 | M-027533-01 |
| ACTB | 60 | M-003451-03 | ATP5E | 514 | M-012330-00 | C1orf41 | 51668 | M-010908-01 |
| ACTC1 | 70 | M-012015-01 | ATP5F1 | 515 | M-015956-01 | C1orf78 | 55194 | M-021255-00 |
| ACTG1 | 71 | M-005265-01 | ATP5G1 | 516 | M-019935-01 | C1QBP | 708 | M-011225-00 |
| ACTN2 | 88 | M-011196-02 | ATP5G2 | 517 | M-019936-01 | C1QR1 | 22918 | M-007815-02 |
| ADCK4 | 79934 | M-005305-01 | ATP5G3 | 518 | M-019494-01 | C20ORF155 | 54675 | M-017910-00 |
| ADORA2A | 135 | M-005416-02 | ATP5H | 10476 | M-012733-02 | C20ORF52 | 140823 | M-015268-02 |
| ADRBK1 | 156 | M-004325-02 | ATP5I | 521 | M-019688-01 | C20ORF7 | 79133 | M-014317-02 |
| AFG3L2 | 10939 | M-005781-00 | ATP5J | 522 | M-017536-01 | C21ORF2 | 755 | M-019913-01 |
| AGER | 177 | M-003625-02 | ATP5J2 | 9551 | M-012690-02 | C22orf29 | 79680 | M-016338-01 |
| AGPAT5 | 55326 | M-008554-02 | ATP5L | 10632 | M-017969-01 | C22orf32 | 91689 | M-016704-00 |
| AGTR2 | 186 | M-005429-02 | ATP5O | 539 | M-019495-00 | C2ORF18 | 54978 | M-018201-01 |
| AHR | 196 | M-004990-01 | ATP5S | 27109 | M-020544-02 | C2orf33 | 56947 | M-018261-01 |
| AK2 | 204 | M-006812-00 | ATP7A | 538 | M-019280-01 | C2orf64 | 493753 | M-034899-00 |
| AKAP1 | 8165 | M-011426-02 | ATPIF1 | 93974 | M-017220-00 | C3 | 718 | M-011001-02 |
| AKAP10 | 11216 | M-017300-01 | AUH | 549 | M-008457-00 | C3ORF1 | 51300 | M-021164-01 |
| AKIP | 54998 | M-006467-00 | BAD | 572 | M-003870-02 | C3orf28 | 26355 | M-016642-02 |
| AKR1B10 | 57016 | M-009691-01 | BAG1 | 573 | M-003871-02 | C3orf31 | 132001 | M-016534-01 |
| AKT2 | 208 | M-003001-02 | BAG3 | 9531 | M-011957-01 | C3orf60 | 25915 | M-027327-01 |
| AKT3 | 10000 | M-003002-02 | BAG5 | 9529 | M-011960-01 | C4ORF14 | 84273 | M-014851-01 |
| ALAS2 | 212 | M-008589-01 | BAK1 | 578 | M-003305-02 | C4orf35 | 85438 | M-018644-01 |
| ALDH1B1 | 219 | M-008254-00 | BAX | 581 | M-003308-03 | C5 | 727 | M-007819-00 |
| ALDH3A2 | 224 | M-009386-03 | BBC3 | 27113 | M-004380-01 | C6ORF125 | 84300 | M-021290-01 |
| ALKBH7 | 84266 | M-014847-00 | BBOX1 | 8424 | M-011472-01 | C6ORF168 | 84553 | M-014910-00 |
| ALOX12 | 239 | M-004558-01 | BCAP31 | 10134 | M-018679-00 | C6ORF49 | 29964 | M-006979-01 |
| ALOX5 | 240 | M-004530-02 | BCDO2 | 83875 | M-014757-01 | C6ORF66 | 29078 | M-020684-01 |
| ALS2CR3 | 66008 | M-014141-00 | BCKDHB | 594 | M-009464-03 | C7ORF23 | 79161 | M-004156-00 |
| AMBRA1 | 55626 | M-029987-01 | BCL2 | 596 | M-003307-06 | C7ORF27 | 221927 | M-016785-01 |
| AMFR | 267 | M-006522-01 | BCL2A1 | 597 | M-003306-01 | C7orf44 | 55744 | M-020238-01 |
| AMID | 84883 | M-004443-01 | BCL2L1 | 598 | M-003458-06 | C8orf37 | 157657 | M-017967-00 |
| ANKRD15 | 23189 | M-012879-01 | BCL2L10 | 10017 | M-004382-01 | C8orf38 | 137682 | M-016179-01 |

#### Table S1

| | | | | | | | | |
|---|---|---|---|---|---|---|---|---|
| **ANXA1** | 301 | M-011161-01 | **BCL2L11** | 10018 | M-004383-02 | **C9ORF111** | 375775 | M-010312-01 |
| **ANXA5** | 308 | M-011209-01 | **BCL2L13** | 23786 | M-020290-00 | **C9ORF89** | 84270 | M-016972-01 |
| **ANXA6** | 309 | M-011210-01 | **BCL2L14** | 79370 | M-004385-03 | **C9ORF90** | 203245 | M-021314-01 |
| **CA4** | 762 | M-008775-00 | **CLIC4** | 25932 | M-013553-00 | **CYP1B1** | 1545 | M-008282-00 |
| **CABC1** | 56997 | M-004939-02 | **CLN8** | 2055 | M-013304-01 | **CYP24A1** | 1591 | M-009269-01 |
| **CACNA2D3** | 55799 | M-021246-01 | **CLPP** | 8192 | M-005811-01 | **CYP27A1** | 1593 | M-008233-00 |
| **CAMK2A** | 815 | M-004942-00 | **CLPX** | 10845 | M-008763-00 | **CYP27B1** | 1594 | M-009757-01 |
| **CAMK2G** | 818 | M-004536-01 | **CLRN1** | 7401 | M-021440-00 | **CYP2E1** | 1571 | M-010134-01 |
| **CAMLG** | 819 | M-011601-01 | **CLU** | 1191 | M-019513-00 | **CYP3A4** | 1576 | M-008169-00 |
| **CAPN1** | 823 | M-005799-00 | **CNGA1** | 1259 | M-006158-01 | **CYP3A5** | 1577 | M-009684-00 |
| **CAPS** | 828 | M-011823-00 | **CNGA3** | 1261 | M-006159-02 | **D1S155E** | 7812 | M-015834-01 |
| **CASP1** | 834 | M-004401-03 | **CNGB1** | 1258 | M-006160-01 | **DACH1** | 1602 | M-013222-01 |
| **CASP2** | 835 | M-003465-03 | **CNP** | 1267 | M-018646-01 | **DAO** | 1610 | M-009756-02 |
| **CASP3** | 836 | M-004307-02 | **CNR2** | 1269 | M-005469-01 | **DAOA** | 267012 | M-015665-01 |
| **CASP6** | 839 | M-004406-02 | **COASY** | 80347 | M-006751-00 | **DAP13** | 55967 | M-009094-00 |
| **CASP7** | 840 | M-004407-02 | **COCH** | 1690 | M-011845-00 | **DAP3** | 7818 | M-004416-00 |
| **CASP8** | 841 | M-003466-05 | **COQ10A** | 93058 | M-018734-01 | **DAPK1** | 1612 | M-004417-03 |
| **CASP9** | 842 | M-003309-01 | **COQ10B** | 80219 | M-018559-01 | **DCI** | 1632 | M-008794-01 |
| **CASQ1** | 844 | M-011227-01 | **COQ2** | 27235 | M-018361-01 | **DCN** | 1634 | M-021491-00 |
| **CAT** | 847 | M-010021-01 | **COQ3** | 51805 | M-010155-01 | **DDHD1** | 80821 | M-021963-01 |
| **CBARA1** | 10367 | M-012720-01 | **COQ4** | 51117 | M-021015-01 | **DDX1** | 1653 | M-011993-00 |
| **CBFA2T3** | 863 | M-017195-00 | **COQ6** | 51004 | M-009942-02 | **DDX17** | 10521 | M-013450-01 |
| **CBLB** | 868 | M-003004-02 | **COQ7** | 10229 | M-017296-00 | **DDX18** | 8886 | M-013451-00 |
| **CCDC109A** | 90550 | M-015519-00 | **COX10** | 1352 | M-013022-00 | **DDX19** | 11269 | M-013471-01 |
| **CCDC109B** | 55013 | M-016108-00 | **COX11** | 1353 | M-011837-00 | **DDX20** | 11218 | M-013472-00 |
| **CCDC123** | 84902 | M-021334-01 | **COX15** | 1355 | M-021442-01 | **DDX21** | 9188 | M-011919-01 |
| **CCDC50** | 152137 | M-017781-01 | **COX18** | 285521 | M-018117-01 | **DDX23** | 9416 | M-019861-00 |
| **CCDC56** | 28958 | M-020558-01 | **COX4I1** | 1327 | M-011625-00 | **DDX24** | 57062 | M-010397-01 |
| **CCDC59** | 29080 | M-020693-01 | **COX4I2** | 84701 | M-013590-00 | **DDX31** | 64794 | M-012931-01 |
| **CCDC90A** | 63933 | M-010730-01 | **COX5A** | 9377 | M-011940-00 | **DDX39** | 10212 | M-004920-01 |
| **CCDC90B** | 60492 | M-016774-00 | **COX5B** | 1329 | M-013632-00 | **DDX3X** | 1654 | M-006874-01 |
| **CCNB1** | 891 | M-003206-02 | **COX6A1** | 1337 | M-011836-00 | **DDX41** | 51428 | M-010394-00 |
| **CCS** | 9973 | M-008954-01 | **COX6B1** | 1340 | M-013148-00 | **DDX42** | 11325 | M-012393-00 |
| **CD163L1** | 283316 | M-008024-02 | **COX6B2** | 125965 | M-019413-02 | **DDX47** | 51202 | M-016299-00 |
| **CD34** | 947 | M-019503-01 | **COX6C** | 1345 | M-013151-01 | **DDX48** | 9775 | M-020762-00 |
| **CD36** | 948 | M-010206-01 | **COX7A1** | 1346 | M-013152-00 | **DDX5** | 1655 | M-003774-01 |
| **CD68** | 968 | M-011236-01 | **COX7A2** | 1347 | M-011626-02 | **DDX50** | 79009 | M-004255-00 |
| **CDC2** | 983 | M-003224-03 | **COX7A2L** | 9167 | M-013179-00 | **DDX54** | 79039 | M-017128-01 |
| **CDH23** | 64072 | M-013051-01 | **COX7B** | 1349 | M-011627-00 | **DDX55** | 57696 | M-027082-01 |
| **CDK2** | 1017 | M-003236-04 | **COX7B2** | 170712 | M-015309-01 | **DDX56** | 54606 | M-020410-01 |
| **CDK5** | 1020 | M-003239-01 | **COX7C** | 1350 | M-013317-00 | **DDX58** | 23586 | M-012511-01 |
| **CDK9** | 1025 | M-003243-03 | **COX8A** | 1351 | M-011819-01 | **DES** | 1674 | M-011638-02 |
| **CDKN1C** | 1028 | M-003244-03 | **COX8C** | 341947 | M-019415-01 | **DFNB31** | 25861 | M-026195-01 |
| **CDS1** | 1040 | M-008680-01 | **CPOX** | 1371 | M-015972-00 | **DHDDS** | 79947 | M-010399-01 |
| **CDS2** | 8760 | M-009591-01 | **CPS1** | 1373 | M-009275-01 | **DHFRL1** | 200895 | M-009013-03 |
| **CDW92** | 23446 | M-010708-01 | **CPT1A** | 1374 | M-009749-02 | **DHODH** | 1723 | M-009619-00 |
| **CENTA2** | 55803 | M-020444-02 | **CPT1B** | 1375 | M-010266-01 | **DHRS1** | 115817 | M-008577-00 |
| **CFH** | 3075 | M-007920-02 | **CPT1C** | 126129 | M-008824-01 | **DHX34** | 9704 | M-032233-01 |
| **CFLAR** | 8837 | M-003772-06 | **CPT2** | 1376 | M-008574-01 | **DHX38** | 9785 | M-013428-01 |
| **CHCHD1** | 118487 | M-027351-00 | **CRAT** | 1384 | M-009524-00 | **DIA1** | 1727 | M-009554-01 |
| **CHCHD3** | 54927 | M-020803-01 | **CRB1** | 23418 | M-012404-01 | **DIABLO** | 56616 | M-004447-00 |
| **CHCHD4** | 131474 | M-016042-01 | **CRTC1** | 23373 | M-014026-02 | **DIAPH1** | 1729 | M-010347-02 |
| **CHCHD6** | 84303 | M-014866-00 | **CRTC2** | 200186 | M-018947-02 | **DIP** | 23151 | M-024634-01 |
| **CHCHD7** | 79145 | M-014326-01 | **CS** | 1431 | M-009334-01 | **DISC1** | 27185 | M-020321-02 |
| **CHDH** | 55349 | M-008123-01 | **CSNK2A1** | 1457 | M-003475-03 | **DKFZP566** | 81889 | M-014695-01 |
| **CHPF** | 79586 | M-017704-00 | **CSPG5** | 10675 | M-020164-01 | **DLAT** | 1737 | M-008490-00 |
| **CHST3** | 9469 | M-003958-01 | **CTSB** | 1508 | M-004266-03 | **DLD** | 1738 | M-009509-01 |
| **CHUK** | 1147 | M-003473-02 | **CTSD** | 1509 | M-003649-00 | **DLK1** | 8788 | M-015911-01 |
| **CIAS1** | 114548 | M-017367-00 | **CUTA** | 51596 | M-020978-01 | **DLST** | 1743 | M-009941-02 |
| **CIG5** | 91543 | M-015423-01 | **CYB5** | 1528 | M-019621-01 | **DMPK** | 1760 | M-004637-01 |
| **CISD1** | 55847 | M-020954-01 | **CYB5B** | 80777 | M-014633-01 | **DNAJA2** | 10294 | M-012104-01 |
| **CISD2** | 493856 | M-032593-00 | **CYBB** | 1536 | M-011021-01 | **DNAJA3** | 9093 | M-017792-00 |

**Table S1**

| | | | | | | | | |
|---|---|---|---|---|---|---|---|---|
| **CKMT1A** | 548596 | M-034935-01 | **CYC1** | 1537 | M-016697-01 | **DNAJC11** | 55735 | M-021205-01 |
| **CKMT1B** | 1159 | M-006708-01 | **CYCS** | 54205 | M-017355-00 | **DNAJC15** | 29103 | M-020286-01 |
| **CKMT2** | 1160 | M-006709-01 | **CYP11A1** | 1583 | M-008329-02 | **DNAJC19** | 131118 | M-016024-00 |
| **CLCN3** | 1182 | M-006151-02 | **CYP11B1** | 1584 | M-008635-02 | **DNCL1** | 8655 | M-005281-02 |
| **CLDN1** | 9076 | M-017369-01 | **CYP11B2** | 1585 | M-009021-01 | **DNLZ** | 728489 | M-184311-01 |
| **CLIC1** | 1192 | M-009530-00 | **CYP1A1** | 1543 | M-004790-02 | **DNM1L** | 10059 | M-012092-01 |
| **DNMT1** | 1786 | M-004605-01 | **FUNDC1** | 139341 | M-018480-01 | **HIGD2A** | 192286 | M-016491-01 |
| **DSG2** | 1829 | M-011645-01 | **FUNDC2** | 65991 | M-006628-01 | **HIP1** | 3092 | M-005001-01 |
| **DUSP1** | 1843 | M-003484-02 | **FXC1** | 26515 | M-018242-02 | **HK1** | 3098 | M-006820-01 |
| **DUSP21** | 63904 | M-007893-01 | **GABARAPL1** | 23710 | M-014715-01 | **HK2** | 3099 | M-006735-01 |
| **DYNC1H1** | 1778 | M-006828-02 | **GABRA5** | 2558 | M-006166-00 | **HK3** | 3101 | M-006736-00 |
| **DYNLL2** | 140735 | M-006493-01 | **GATA1** | 2623 | M-009610-00 | **HMGB1** | 3146 | M-018981-01 |
| **DYNLT1** | 6993 | M-019964-00 | **GATM** | 2628 | M-008900-01 | **HMGCL** | 3155 | M-019290-01 |
| **E2F1** | 1869 | M-003259-01 | **GCDH** | 2639 | M-004039-01 | **HMGCS1** | 3157 | M-009808-01 |
| **ECGF1** | 1890 | M-009281-01 | **GCH1** | 2643 | M-010328-03 | **HMGCS2** | 3158 | M-010179-01 |
| **ECH1** | 1891 | M-004035-01 | **GDAP1** | 54332 | M-021225-01 | **HMOX1** | 3162 | M-006372-02 |
| **ECHS1** | 1892 | M-010343-00 | **GDF5** | 8200 | M-012271-02 | **HNRPM** | 4670 | M-013452-01 |
| **EFEMP1** | 2202 | M-011855-01 | **GDNF** | 2668 | M-011040-01 | **HRAS** | 3265 | M-004142-00 |
| **EFHA2** | 286097 | M-018623-00 | **GFAP** | 2670 | M-011667-00 | **HSA9947** | 23400 | M-008601-01 |
| **EFHD1** | 80303 | M-010673-00 | **GFER** | 2671 | M-012041-01 | **HSCB** | 150274 | M-017718-01 |
| **EGFL11** | 346007 | M-024856-00 | **GGA2** | 23062 | M-012908-01 | **HSD17B8** | 7923 | M-008141-02 |
| **EGFR** | 1956 | M-003114-03 | **GGT1** | 2678 | M-005884-01 | **HSD3B1** | 3283 | M-008972-02 |
| **EI24** | 9538 | M-019879-01 | **GHITM** | 27069 | M-020534-01 | **HSD3B2** | 3284 | M-012542-01 |
| **EIF2S1** | 1965 | M-015389-01 | **GIMAP5** | 55340 | M-013342-01 | **HSP90B1** | 7184 | M-006417-02 |
| **EIF3S10** | 8661 | M-019534-01 | **GJA1** | 2697 | M-011042-01 | **HSPA1A** | 3303 | M-005168-01 |
| **EIF4G1** | 1981 | M-019474-01 | **GJB2** | 2706 | M-019285-01 | **HSPA1B** | 3304 | M-003501-03 |
| **EIF4G2** | 1982 | M-011263-01 | **GJB3** | 2707 | M-019948-02 | **HSPA4** | 3308 | M-012636-02 |
| **ELA2** | 1991 | M-005861-02 | **GJB6** | 10804 | M-019916-01 | **HSPA5** | 3309 | M-008198-02 |
| **ENDOGL1** | 9941 | M-008572-01 | **GK** | 2710 | M-006727-00 | **HSPA9B** | 3313 | M-004750-03 |
| **ENOSF1** | 55556 | M-020894-00 | **GK2** | 2712 | M-015091-01 | **HSPB1** | 3315 | M-005269-01 |
| **ENOX1** | 55068 | M-021077-00 | **GLRX** | 2745 | M-012634-01 | **HSPB8** | 26353 | M-005006-00 |
| **EPHA4** | 2043 | M-003118-02 | **GLRX2** | 51022 | M-021332-01 | **HSPCA** | 3320 | M-005186-02 |
| **EPHB6** | 2051 | M-003125-02 | **GLTSCR2** | 29997 | M-006464-01 | **HSPD1** | 3329 | M-010600-02 |
| **EPO** | 2056 | M-020204-01 | **GMIP** | 51291 | M-021160-01 | **HSPE1** | 3336 | M-019649-00 |
| **ERAL1** | 26284 | M-012709-01 | **GNA12** | 2768 | M-008435-00 | **IAPP** | 3375 | M-007790-01 |
| **ERBB4** | 2066 | M-003128-03 | **GNB2L1** | 10399 | M-006876-01 | **IBRDC2** | 255488 | M-025119-01 |
| **ERCC6** | 2074 | M-004888-01 | **GOLPH3** | 64083 | M-006414-00 | **ICT1** | 3396 | M-010517-00 |
| **ETFA** | 2108 | M-011029-01 | **GOT2** | 2806 | M-011674-02 | **IDE** | 3416 | M-005899-03 |
| **ETFB** | 2109 | M-010494-01 | **GPAM** | 57678 | M-009946-01 | **IDH2** | 3418 | M-004013-00 |
| **ETFDH** | 2110 | M-008127-02 | **GPD2** | 2820 | M-009843-02 | **IER3** | 8870 | M-011547-01 |
| **EYA2** | 2139 | M-017233-01 | **GPR30** | 2852 | M-005563-02 | **IF** | 3426 | M-005900-02 |
| **F2R** | 2149 | M-005094-01 | **GPR81** | 27198 | M-005601-02 | **IFI27** | 3429 | M-006465-02 |
| **FAM36A** | 116228 | M-021429-01 | **GPX4** | 2879 | M-011676-01 | **IFI6** | 2537 | M-003672-02 |
| **FAM82C** | 55177 | M-020973-00 | **GRHL2** | 79977 | M-014515-01 | **IFNA2** | 3440 | M-013809-01 |
| **FARSLB** | 10056 | M-015414-01 | **GRP58** | 2923 | M-003674-01 | **IFNB1** | 3456 | M-019656-01 |
| **FASLG** | 356 | M-011130-00 | **GRPEL2** | 134266 | M-016191-01 | **IGF1R** | 3480 | M-003012-05 |
| **FATE1** | 89885 | M-015068-01 | **GSK3A** | 2931 | M-003009-01 | **IGF2** | 3481 | M-004093-01 |
| **FCGR2B** | 2213 | M-015823-01 | **GSK3B** | 2932 | M-003010-03 | **IKBKB** | 3551 | M-003503-03 |
| **FCGR3B** | 2215 | M-019374-01 | **GSN** | 2934 | M-007775-03 | **IKBKE** | 9641 | M-003723-02 |
| **FECH** | 2235 | M-011036-01 | **GSR** | 2936 | M-009647-01 | **IL7** | 3574 | M-007995-01 |
| **FEN1** | 2237 | M-010344-01 | **GSTA4** | 2941 | M-011289-00 | **IMMP2L** | 83943 | M-005902-02 |
| **FGFBP1** | 9982 | M-019910-00 | **GSTK1** | 373156 | M-020958-01 | **IMMT** | 10989 | M-019832-01 |
| **FGR** | 2268 | M-003135-03 | **GTPBP5** | 26164 | M-013036-01 | **IMPG2** | 50939 | M-020890-01 |
| **FHIT** | 2272 | M-004952-02 | **GUCA1B** | 2979 | M-015131-01 | **INS** | 3630 | M-011058-01 |
| **FIS1** | 51024 | M-020907-02 | **GUF1** | 60558 | M-021270-01 | **INSIG2** | 51141 | M-021039-00 |
| **FKBP5** | 2289 | M-004224-01 | **GZMB** | 3002 | M-005889-02 | **IPLA2gamma** | 50640 | M-010284-01 |
| **FKBP8** | 23770 | M-009673-02 | **GZMH** | 2999 | M-005890-02 | **IRF1** | 3659 | M-011704-01 |
| **FKSG24** | 84769 | M-014959-02 | **HAAO** | 23498 | M-008666-00 | **IRS1** | 3667 | M-003015-01 |
| **FKTN** | 2218 | M-012313-01 | **HADH2** | 3028 | M-009390-01 | **ITGA11** | 22801 | M-008000-02 |
| **FLCN** | 201163 | M-009998-01 | **HADHA** | 3030 | M-009470-01 | **ITGA2** | 3673 | M-004566-02 |
| **FLJ12592** | 84129 | M-008129-01 | **HADHB** | 3032 | M-008280-01 | **ITPR3** | 3710 | M-006209-02 |
| **FLJ14466** | 84876 | M-014998-01 | **HADHSC** | 3033 | M-008298-01 | **ITSN1** | 6453 | M-008365-01 |

**Table S1**

| | | | | | | | | |
|---|---|---|---|---|---|---|---|---|
| **FLJ25059** | 196294 | M-005877-01 | **HAX1** | 10456 | M-012168-01 | **JTB** | 10899 | M-010567-01 |
| **FLJ30473** | 150209 | M-008134-00 | **HBB** | 3043 | M-011047-01 | **KARS** | 3735 | M-012114-00 |
| **FLVCR** | 28982 | M-020584-01 | **HCCS** | 3052 | M-009226-01 | **KCNA3** | 3738 | M-006213-01 |
| **FMR1** | 2332 | M-019631-00 | **HD** | 3064 | M-003737-02 | **KCNA5** | 3741 | M-006215-01 |
| **FNDC1** | 84624 | M-024901-01 | **HDAC6** | 10013 | M-003499-00 | **KCNJ10** | 3766 | M-006240-01 |
| **FOXO3A** | 2309 | M-003007-02 | **HEBP2** | 23593 | M-020612-00 | **KCNK3** | 3777 | M-006262-02 |
| **FOXRED1** | 55572 | M-008137-01 | **HERC2** | 8924 | M-007180-02 | **KCNK9** | 51305 | M-004891-01 |
| **FPGS** | 2356 | M-016472-01 | **HGF** | 3082 | M-006650-01 | **KCNN3** | 3782 | M-006270-02 |
| **FTH1** | 2495 | M-019634-02 | **HIGD1A** | 25994 | M-020242-01 | **KCNN4** | 3783 | M-004461-01 |
| **KCNQ4** | 9132 | M-006274-00 | **MCART1** | 92014 | M-007358-01 | **MRPL41** | 64975 | M-013572-00 |
| **KDR** | 3791 | M-003148-01 | **MCART2** | 147407 | M-031039-01 | **MRPL42** | 28977 | M-017553-00 |
| **KIAA0174** | 9798 | M-020977-00 | **MCART6** | 401612 | M-031566-02 | **MRPL43** | 84545 | M-019126-00 |
| **KIAA0446** | 9673 | M-007339-01 | **MCCC1** | 56922 | M-009429-02 | **MRPL44** | 65080 | M-012858-00 |
| **KIAA0774** | 23281 | M-006850-01 | **MCL1** | 4170 | M-004501-08 | **MRPL45** | 84311 | M-019256-02 |
| **KIAA0831** | 22863 | M-020438-01 | **MDH2** | 4191 | M-008439-00 | **MRPL46** | 26589 | M-017562-00 |
| **KIF14** | 9928 | M-003319-00 | **ME2** | 4200 | M-009461-01 | **MRPL47** | 57129 | M-013186-01 |
| **KIF1B** | 23095 | M-009317-01 | **MECP2** | 4204 | M-013094-02 | **MRPL48** | 51642 | M-017512-02 |
| **KIF5B** | 3799 | M-008867-00 | **MEF2A** | 4205 | M-009362-00 | **MRPL49** | 740 | M-017507-00 |
| **KLHL7** | 55975 | M-015574-00 | **MEF2C** | 4208 | M-009455-00 | **MRPL50** | 54534 | M-013373-02 |
| **KMO** | 8564 | M-009897-01 | **MEF2D** | 4209 | M-009884-00 | **MRPL51** | 51258 | M-013563-00 |
| **KRT18** | 3875 | M-010604-02 | **MERTK** | 10461 | M-003155-02 | **MRPL52** | 122704 | M-019012-01 |
| **KRT6A** | 3853 | M-012116-01 | **MFN1** | 55669 | M-010670-01 | **MRPL53** | 116540 | M-012941-00 |
| **KRT8** | 3856 | M-019658-01 | **MFN2** | 9927 | M-012961-00 | **MRPL54** | 116541 | M-017497-01 |
| **KUB3** | 91419 | M-015098-01 | **MFTC** | 81034 | M-007356-00 | **MRPL55** | 128308 | M-019054-02 |
| **LAMB2** | 3913 | M-013310-01 | **MGC4767** | 84274 | M-009755-01 | **MRPL9** | 65005 | M-013498-02 |
| **LAMC3** | 10319 | M-012173-01 | **MGC5352** | 192111 | M-015919-02 | **MRPS10** | 55173 | M-013345-00 |
| **LASS6** | 253782 | M-032207-00 | **MGEA5** | 10724 | M-012805-01 | **MRPS11** | 64963 | M-013623-00 |
| **LCK** | 3932 | M-003151-02 | **MGST1** | 4257 | M-009248-00 | **MRPS12** | 6183 | M-011382-01 |
| **LDB3** | 11155 | M-026288-02 | **MIPEP** | 4285 | M-005949-02 | **MRPS14** | 63931 | M-013050-00 |
| **LDHA** | 3939 | M-008201-01 | **MJD** | 4287 | M-012013-01 | **MRPS15** | 64960 | M-013609-00 |
| **LDHB** | 3945 | M-009779-01 | **MKKS** | 8195 | M-013300-00 | **MRPS16** | 51021 | M-013130-01 |
| **LDHD** | 197257 | M-008761-01 | **MK-STYX** | 51657 | M-008031-01 | **MRPS17** | 51373 | M-013215-00 |
| **LETM1** | 3954 | M-019549-00 | **MLXIP** | 22877 | M-008976-01 | **MRPS18A** | 55168 | M-007008-00 |
| **LETM2** | 137994 | M-015974-01 | **MMAA** | 166785 | M-016320-01 | **MRPS18B** | 28973 | M-013043-00 |
| **LETMD1** | 25875 | M-016920-01 | **MMP1** | 4312 | M-005951-01 | **MRPS18C** | 51023 | M-013131-00 |
| **LGALS3** | 3958 | M-010606-02 | **MOAP1** | 64112 | M-004430-02 | **MRPS21** | 54460 | M-013388-00 |
| **LHFPL5** | 222662 | M-018911-01 | **MOSC1** | 64757 | M-019358-00 | **MRPS22** | 56945 | M-013219-00 |
| **LMNA** | 4000 | M-004978-01 | **MOSC2** | 54996 | M-018689-00 | **MRPS23** | 51649 | M-012973-01 |
| **LOC150763** | 150763 | M-010302-01 | **MPST** | 4357 | M-010119-01 | **MRPS24** | 64951 | M-013534-02 |
| **LOC153328** | 153328 | M-007347-02 | **MPV17** | 4358 | M-017720-01 | **MRPS25** | 64432 | M-010142-00 |
| **LOC201164** | 201164 | M-017858-00 | **MPV17L** | 255027 | M-018370-00 | **MRPS26** | 64949 | M-013546-00 |
| **LOC203427** | 203427 | M-007349-01 | **MRC1** | 4360 | M-011730-01 | **MRPS27** | 23107 | M-012903-00 |
| **LOC283130** | 283130 | M-007351-01 | **M-RIP** | 23164 | M-014102-01 | **MRPS28** | 28957 | M-013492-00 |
| **LOC790955** | 790955 | M-184603-01 | **MRPL1** | 65008 | M-017264-01 | **MRPS30** | 10884 | M-013207-02 |
| **LPIN1** | 23175 | M-017427-01 | **MRPL10** | 124995 | M-017394-00 | **MRPS31** | 10240 | M-012111-00 |
| **LPL** | 4023 | M-008970-01 | **MRPL11** | 65003 | M-013123-00 | **MRPS33** | 51650 | M-012940-00 |
| **LRAT** | 9227 | M-010272-01 | **MRPL12** | 6182 | M-017517-00 | **MRPS34** | 65993 | M-012884-00 |
| **LRPPRC** | 10128 | M-018773-00 | **MRPL13** | 28998 | M-013557-00 | **MRPS35** | 60488 | M-013071-00 |
| **LRRC51** | 220074 | M-016302-01 | **MRPL14** | 64928 | M-013550-01 | **MRPS36** | 92259 | M-019052-01 |
| **LRRC59** | 55379 | M-010669-00 | **MRPL15** | 29088 | M-013052-00 | **MRPS5** | 64969 | M-019247-00 |
| **LRRK2** | 120892 | M-006323-02 | **MRPL16** | 54948 | M-013354-00 | **MRPS6** | 64968 | M-019243-00 |
| **LYN** | 4067 | M-003153-04 | **MRPL17** | 63875 | M-013049-00 | **MRPS7** | 51081 | M-013580-00 |
| **LYRM7** | 90624 | M-018887-01 | **MRPL18** | 29074 | M-017251-00 | **MRPS9** | 64965 | M-019184-00 |
| **MAG** | 4099 | M-011722-00 | **MRPL19** | 9801 | M-013418-01 | **MRS2L** | 57380 | M-020748-02 |
| **MAGMAS** | 51025 | M-015261-00 | **MRPL2** | 51069 | M-017261-00 | **MSRA** | 4482 | M-012464-00 |
| **MAOA** | 4128 | M-009369-01 | **MRPL20** | 55052 | M-017564-00 | **MSTO1** | 55154 | M-021170-00 |
| **MAOB** | 4129 | M-010183-03 | **MRPL21** | 219927 | M-019072-02 | **MT2A** | 4502 | M-018338-00 |
| **MAP1A** | 4130 | M-013482-01 | **MRPL22** | 29093 | M-017259-01 | **MTCH1** | 23787 | M-007370-01 |
| **MAP1LC3A** | 84557 | M-013579-00 | **MRPL23** | 6150 | M-013124-00 | **MTCH2** | 23788 | M-007371-00 |
| **MAP1LC3B** | 81631 | M-012846-01 | **MRPL24** | 79590 | M-017442-00 | **MTFR1** | 9650 | M-019432-00 |
| **MAP2K1** | 5604 | M-003571-01 | **MRPL27** | 51264 | M-013182-01 | **MTG1** | 92170 | M-015522-01 |
| **MAP2K2** | 5605 | M-003573-03 | **MRPL28** | 10573 | M-017627-00 | **MTHFD1L** | 25902 | M-009949-01 |

#### Table S1

| | | | | | | | | |
|---|---|---|---|---|---|---|---|---|
| **MAP2K3** | 5606 | M-003509-03 | **MRPL3** | 11222 | M-012372-00 | **MTHFD2L** | 441024 | M-032402-01 |
| **MAP3K12** | 7786 | M-003312-02 | **MRPL30** | 51263 | M-013181-01 | **MTNR1A** | 4543 | M-005669-02 |
| **MAPK12** | 6300 | M-003590-03 | **MRPL32** | 64983 | M-013512-01 | **MTP18** | 51537 | M-021196-01 |
| **MAPK3** | 5595 | M-003592-03 | **MRPL33** | 9553 | M-017401-01 | **MTRF1** | 9617 | M-019211-00 |
| **MAPK8** | 5599 | M-003514-04 | **MRPL34** | 64981 | M-012885-00 | **MTUS1** | 57509 | M-006848-00 |
| **MAPK8IP1** | 9479 | M-003595-00 | **MRPL35** | 51318 | M-013206-01 | **MTX1** | 4580 | M-019667-02 |
| **MAPKAP1** | 79109 | M-014315-02 | **MRPL36** | 64979 | M-013574-00 | **MTX2** | 10651 | M-020087-01 |
| **MAPRE1** | 22919 | M-006824-00 | **MRPL37** | 51253 | M-017460-00 | **MTX3** | 345778 | M-024528-01 |
| **MARCH5** | 54708 | M-007001-01 | **MRPL38** | 64978 | M-013573-00 | **MUC1** | 4582 | M-004019-02 |
| **MARK2** | 2011 | M-004260-02 | **MRPL39** | 54148 | M-008542-01 | **MULK** | 55750 | M-007256-02 |
| **MARVELD2** | 153562 | M-017054-02 | **MRPL4** | 51073 | M-017470-00 | **MYH11** | 4629 | M-011737-01 |
| **MB** | 4151 | M-012057-01 | **MRPL40** | 64976 | M-017608-00 | **MYH14** | 79784 | M-027149-01 |
| **MYH6** | 4624 | M-012645-01 | **NNT** | 23530 | M-009809-01 | **PID1** | 55022 | M-018934-01 |
| **MYH9** | 4627 | M-007668-01 | **NOS1** | 4842 | M-009496-01 | **PIK3CA** | 5290 | M-003018-03 |
| **MYLK** | 4638 | M-005351-05 | **NOTCH1** | 4851 | M-007771-02 | **PIK3CG** | 5294 | M-005274-02 |
| **MYO19** | 80179 | M-017137-02 | **NOTCH3** | 4854 | M-011093-01 | **PIK4CB** | 5298 | M-006777-03 |
| **MYO1A** | 4640 | M-008765-00 | **NOX1** | 27035 | M-010193-01 | **PIM1** | 5292 | M-003923-00 |
| **MYO6** | 4646 | M-006355-00 | **NOX4** | 50507 | M-010194-00 | **PINK1** | 65018 | M-004030-02 |
| **MYO7A** | 4647 | M-019330-01 | **NPC1** | 4864 | M-008047-01 | **PISD** | 23761 | M-009548-00 |
| **MYOC** | 4653 | M-011089-02 | **NPTX1** | 4884 | M-011343-01 | **PLA2G2A** | 5320 | M-009901-03 |
| **NAT8L** | 339983 | M-009115-00 | **NR2C2** | 7182 | M-003418-02 | **PLA2G4A** | 5321 | M-009886-01 |
| **NBL1** | 4681 | M-006540-01 | **NR4A1** | 3164 | M-003426-03 | **PLA2G6** | 8398 | M-009085-04 |
| **NCF1** | 653361 | M-180696-00 | **NRG1** | 3084 | M-004608-02 | **PLA2G7** | 7941 | M-004903-01 |
| **NCOA7** | 135112 | M-018862-00 | **NTN1** | 9423 | M-011946-01 | **PLAA** | 9373 | M-016215-01 |
| **NDE1** | 54820 | M-020625-00 | **NTRK1** | 4914 | M-003159-02 | **PLAUR** | 5329 | M-006388-01 |
| **NDUFA1** | 4694 | M-011881-01 | **NUBPL** | 80224 | M-021287-01 | **PLB1** | 151056 | M-008684-00 |
| **NDUFA10** | 4705 | M-006752-00 | **NUP93** | 9688 | M-020767-00 | **PLEKHF1** | 79156 | M-018423-01 |
| **NDUFA11** | 126328 | M-018508-00 | **OCIAD2** | 132299 | M-015633-01 | **PLEKHF2** | 79666 | M-018407-00 |
| **NDUFA12L** | 91942 | M-017758-01 | **OGDH** | 4967 | M-009679-02 | **PLG** | 5340 | M-006001-02 |
| **NDUFA13** | 51079 | M-016921-01 | **OGG1** | 4968 | M-005147-03 | **PLN** | 5350 | M-011754-00 |
| **NDUFA2** | 4695 | M-018869-01 | **OLA1** | 29789 | M-015680-01 | **PLOD2** | 5352 | M-004285-01 |
| **NDUFA4** | 4697 | M-019200-00 | **OMA1** | 115209 | M-008662-03 | **PLSCR3** | 57048 | M-010255-00 |
| **NDUFA5** | 4698 | M-012000-00 | **OPA1** | 4976 | M-005273-00 | **PMAIP1** | 5366 | M-005275-03 |
| **NDUFA6** | 4700 | M-015716-01 | **OPA3** | 80207 | M-014595-01 | **PMPCA** | 23203 | M-008734-01 |
| **NDUFA7** | 4701 | M-012693-01 | **OPRD1** | 4985 | M-005683-01 | **PMPCB** | 9512 | M-004747-01 |
| **NDUFA8** | 4702 | M-012496-00 | **OPTN** | 10133 | M-016269-02 | **PNKP** | 11284 | M-006783-02 |
| **NDUFA9** | 4704 | M-016044-02 | **OSAP** | 84709 | M-010682-02 | **PNPLA2** | 57104 | M-009003-01 |
| **NDUFAB1** | 4706 | M-019897-01 | **OTC** | 5009 | M-009291-02 | **PNPLA3** | 80339 | M-009564-01 |
| **NDUFAF1** | 51103 | M-021003-02 | **OTOA** | 146183 | M-016394-01 | **PNPT1** | 87178 | M-019454-00 |
| **NDUFB1** | 4707 | M-017848-02 | **OTOF** | 9381 | M-011942-00 | **POLG** | 5428 | M-012649-00 |
| **NDUFB10** | 4716 | M-012675-01 | **OXA1L** | 5018 | M-012696-00 | **POLR2B** | 5431 | M-011187-00 |
| **NDUFB11** | 54539 | M-016098-02 | **P2RX5** | 5026 | M-006286-04 | **PON2** | 5445 | M-009676-01 |
| **NDUFB2** | 4708 | M-019202-02 | **P2RX7** | 5027 | M-003728-01 | **PON3** | 5446 | M-009675-02 |
| **NDUFB3** | 4709 | M-019604-01 | **PAEP** | 5047 | M-010027-02 | **PP591** | 80308 | M-008629-00 |
| **NDUFB4** | 4710 | M-032508-00 | **PAK1** | 5058 | M-003521-04 | **PPARA** | 5465 | M-003434-01 |
| **NDUFB5** | 4711 | M-019209-02 | **PANK2** | 80025 | M-003797-03 | **PPIF** | 10105 | M-009708-00 |
| **NDUFB6** | 4712 | M-017210-00 | **PAPD1** | 55149 | M-016486-00 | **PPOX** | 5498 | M-008383-01 |
| **NDUFB7** | 4713 | M-017213-01 | **PARG** | 8505 | M-011488-02 | **PPP1CA** | 5499 | M-008927-01 |
| **NDUFB8** | 4714 | M-019898-01 | **PARK2** | 5071 | M-003603-00 | **PPP1R15A** | 23645 | M-004442-01 |
| **NDUFB9** | 4715 | M-019899-01 | **PARK7** | 11315 | M-005984-00 | **PPP2R2B** | 5521 | M-003022-02 |
| **NDUFC1** | 4717 | M-019601-02 | **PARL** | 55486 | M-021387-01 | **PPP3CC** | 5533 | M-010005-00 |
| **NDUFC2** | 4718 | M-015319-01 | **PARP1** | 142 | M-006656-01 | **PPP3R1** | 5534 | M-009869-02 |
| **NDUFS1** | 4719 | M-019069-00 | **PARP4** | 143 | M-007244-03 | **PRCD** | 768206 | M-183754-01 |
| **NDUFS2** | 4720 | M-015770-01 | **PBEF1** | 10135 | M-004581-01 | **PRDX3** | 10935 | M-010355-00 |
| **NDUFS3** | 4722 | M-019815-01 | **PC** | 5091 | M-008950-02 | **PRDX5** | 25824 | M-019102-00 |
| **NDUFS4** | 4724 | M-019602-00 | **PCDH15** | 65217 | M-013654-01 | **PRELID1** | 27166 | M-017650-01 |
| **NDUFS5** | 4725 | M-019816-00 | **PDCD5** | 9141 | M-004439-01 | **PREP** | 5550 | M-006006-01 |
| **NDUFS6** | 4726 | M-019817-01 | **PDCD8** | 9131 | M-011912-00 | **PRKAA1** | 5562 | M-005027-02 |
| **NDUFS7** | 374291 | M-031021-00 | **PDE2A** | 5138 | M-007644-00 | **PRKAB1** | 5564 | M-007675-00 |
| **NDUFS8** | 4728 | M-019600-00 | **PDE6A** | 5145 | M-007651-00 | **PRKACA** | 5566 | M-004649-01 |
| **NDUFV1** | 4723 | M-016266-00 | **PDE6B** | 5158 | M-007652-01 | **PRKAG3** | 53632 | M-009859-01 |
| **NDUFV2** | 4729 | M-012589-01 | **PDE6G** | 5148 | M-007655-01 | **PRKCA** | 5578 | M-003523-03 |

#### Table S1

| | | | | | | | | |
|---|---|---|---|---|---|---|---|---|
| **NDUFV3** | 4731 | M-016362-01 | **PDGFB** | 5155 | M-011749-00 | **PRKCD** | 5580 | M-003524-01 |
| **NF1** | 4763 | M-003916-03 | **PDGFRA** | 5156 | M-003162-04 | **PRKCE** | 5581 | M-004653-02 |
| **NFATC1** | 4772 | M-003605-03 | **PDK4** | 5166 | M-019425-02 | **PRKCM** | 5587 | M-005028-02 |
| **NFE2L2** | 4780 | M-003755-02 | **PDZK1** | 5174 | M-010615-03 | **PRKCQ** | 5588 | M-003525-01 |
| **NFKBIA** | 4792 | M-004765-00 | **PEA15** | 8682 | M-010553-01 | **PRKCZ** | 5590 | M-003526-04 |
| **NFS1** | 9054 | M-011564-00 | **PECI** | 10455 | M-009804-00 | **PRKD2** | 25865 | M-004197-02 |
| **NGFR** | 4804 | M-009340-02 | **PEMT** | 10400 | M-010392-00 | **PRKG1** | 5592 | M-004658-04 |
| **NHEDC2** | 133308 | M-007345-00 | **PEX13** | 5194 | M-012591-02 | **PRKR** | 5610 | M-003527-00 |
| **NIPSNAP1** | 8508 | M-011489-00 | **PEX3** | 8504 | M-019544-00 | **PRNP** | 5621 | M-011101-01 |
| **NLN** | 57486 | M-005977-01 | **PFKP** | 5214 | M-010253-01 | **PRODH** | 5625 | M-009543-00 |
| **NLRX1** | 79671 | M-012926-01 | **PGR** | 5241 | M-003433-01 | **PRODH2** | 58510 | M-013098-01 |
| **NME1** | 4830 | M-006821-01 | **PGS1** | 9489 | M-009483-01 | **PROM1** | 8842 | M-010630-01 |
| **NME2** | 4831 | M-005102-02 | **PHB** | 5245 | M-010530-00 | **PRPF6** | 24148 | M-012821-01 |
| **NME4** | 4833 | M-006494-00 | **PHLDA2** | 7262 | M-011411-01 | **PRPF8** | 10594 | M-012252-02 |
| **NMT1** | 4836 | M-004316-00 | **PI4KII** | 55361 | M-006770-02 | **PRSS11** | 5654 | M-006009-02 |
| **PRSS15** | 9361 | M-003979-00 | **SAC** | 55811 | M-006353-01 | **SLC25A29** | 123096 | M-007318-01 |
| **PRSS25** | 27429 | M-006014-04 | **SAMM50** | 25813 | M-017871-00 | **SLC25A3** | 5250 | M-007484-00 |
| **PSEN1** | 5663 | M-004998-01 | **SARM1** | 23098 | M-008076-01 | **SLC25A30** | 253512 | M-007350-01 |
| **PSEN2** | 5664 | M-006018-02 | **SCN1A** | 6323 | M-006297-02 | **SLC25A31** | 83447 | M-007322-01 |
| **PTCD3** | 55037 | M-016957-00 | **SCN5A** | 6331 | M-006500-03 | **SLC25A33** | 84275 | M-007366-01 |
| **PTEN** | 5728 | M-003023-02 | **SCO1** | 6341 | M-011892-01 | **SLC25A34** | 284723 | M-032041-02 |
| **PTGIS** | 5740 | M-004691-02 | **SCO2** | 9997 | M-011987-01 | **SLC25A35** | 399512 | M-031743-01 |
| **PTK2** | 5747 | M-003164-02 | **SDHA** | 6389 | M-009398-02 | **SLC25A36** | 55186 | M-007327-02 |
| **PTK2B** | 2185 | M-003165-03 | **SDHB** | 6390 | M-011773-02 | **SLC25A37** | 51312 | M-007369-01 |
| **PTMA** | 5757 | M-005207-01 | **SDHC** | 6391 | M-011385-01 | **SLC25A38** | 54977 | M-007331-01 |
| **PTPMT1** | 114971 | M-029988-02 | **SDHD** | 6392 | M-006305-00 | **SLC25A39** | 51629 | M-007319-00 |
| **PTPN1** | 5770 | M-003529-04 | **SEMA4A** | 64218 | M-015686-00 | **SLC25A4** | 291 | M-007485-02 |
| **PTPN11** | 5781 | M-003947-01 | **SERAC1** | 84947 | M-015026-00 | **SLC25A40** | 55972 | M-007354-01 |
| **PTPRC** | 5788 | M-008067-01 | **SERPINA1** | 5265 | M-008847-01 | **SLC25A41** | 284427 | M-007363-01 |
| **PTRH2** | 51651 | M-007271-01 | **SET** | 6418 | M-019586-01 | **SLC25A42** | 284439 | M-007361-01 |
| **PXN** | 5829 | M-005163-00 | **SFN** | 2810 | M-005180-00 | **SLC25A46** | 91137 | M-007353-00 |
| **PYCR1** | 5831 | M-012349-00 | **SFRP4** | 6424 | M-011388-01 | **SLC25A5** | 292 | M-007486-03 |
| **PYCS** | 5832 | M-006785-01 | **SFXN1** | 94081 | M-010686-00 | **SLC25A6** | 293 | M-007487-01 |
| **RAB11A** | 8766 | M-004726-02 | **SFXN2** | 118980 | M-018547-00 | **SLC26A4** | 5172 | M-007493-01 |
| **RAB11FIP1** | 80223 | M-015968-01 | **SFXN3** | 81855 | M-018729-02 | **SLC27A3** | 11000 | M-007499-01 |
| **RAB11FIP5** | 26056 | M-004298-01 | **SFXN4** | 119559 | M-018237-01 | **SLC29A1** | 2030 | M-003709-01 |
| **RAB32** | 10981 | M-009920-02 | **SFXN5** | 94097 | M-016367-01 | **SLC2A1** | 6513 | M-007509-01 |
| **RAC1** | 5879 | M-003560-06 | **SGCD** | 6444 | M-017292-00 | **SLC2A4** | 6517 | M-007517-02 |
| **RAC2** | 5880 | M-007741-01 | **SGK** | 6446 | M-003027-05 | **SLC35B3** | 51000 | M-007544-00 |
| **RAF1** | 5894 | M-003601-02 | **SH120** | 51463 | M-005725-01 | **SLC35C1** | 55343 | M-010693-00 |
| **RALA** | 5898 | M-009235-00 | **SH3BP5** | 9467 | M-019869-01 | **SLC37A4** | 2542 | M-007557-01 |
| **RALBP1** | 10928 | M-009266-00 | **SH3GLB1** | 51100 | M-017086-01 | **SLC38A4** | 55089 | M-007561-01 |
| **RAP1B** | 5908 | M-010364-03 | **SHC1** | 6464 | M-018841-02 | **SLC3A1** | 6519 | M-007575-00 |
| **RDH12** | 145226 | M-008319-00 | **SHH** | 6469 | M-006036-02 | **SLC5A1** | 6523 | M-007589-01 |
| **RDH13** | 112724 | M-008628-00 | **SHMT2** | 6472 | M-004906-01 | **SLC6A1** | 6529 | M-007597-01 |
| **RDS** | 5961 | M-011102-01 | **SIGLEC8** | 27181 | M-012843-01 | **SLC8A2** | 6543 | M-007621-01 |
| **RDX** | 5962 | M-011762-02 | **SIRT1** | 23411 | M-003540-01 | **SLC8A3** | 6547 | M-007622-02 |
| **REA** | 11331 | M-018703-00 | **SIRT3** | 23410 | M-004827-01 | **SLC9A1** | 6548 | M-005277-00 |
| **RECQL4** | 9401 | M-010559-01 | **SIVA** | 10572 | M-012262-02 | **SLC9A3R1** | 9368 | M-012688-01 |
| **REEP1** | 65055 | M-014235-00 | **SLC11A2** | 4891 | M-007381-01 | **SLCO1A2** | 6579 | M-007439-00 |
| **RELA** | 5970 | M-003533-02 | **SLC16A7** | 9194 | M-007409-01 | **SMAD5** | 4090 | M-015791-00 |
| **REN** | 5972 | M-006026-01 | **SLC17A8** | 246213 | M-007418-00 | **SMCP** | 4184 | M-017503-00 |
| **REP15** | 387849 | M-030132-01 | **SLC19A1** | 6573 | M-007422-01 | **SMCR7** | 125170 | M-017272-00 |
| **RET** | 5979 | M-003170-02 | **SLC1A3** | 6507 | M-007427-00 | **SMCR7L** | 54471 | M-015938-00 |
| **RFFL** | 117584 | M-007120-03 | **SLC1A4** | 6509 | M-007428-01 | **SMPD1** | 6609 | M-006676-01 |
| **RGR** | 5995 | M-005721-02 | **SLC22A3** | 6581 | M-007454-00 | **SMPD4** | 55627 | M-020681-02 |
| **RHO** | 6010 | M-005722-00 | **SLC22A7** | 10864 | M-007458-01 | **SMURF1** | 57154 | M-007191-01 |
| **RHOA** | 387 | M-003860-03 | **SLC23A2** | 9962 | M-007462-01 | **SNN** | 8303 | M-018536-01 |
| **RHOT1** | 55288 | M-010365-01 | **SLC24A6** | 80024 | M-007332-00 | **SNPH** | 9751 | M-020417-00 |
| **RHOT2** | 89941 | M-008340-01 | **SLC25A1** | 6576 | M-007468-01 | **SNTA1** | 6640 | M-011777-00 |
| **RIMS3** | 9783 | M-020891-00 | **SLC25A10** | 1468 | M-007469-00 | **SNTG1** | 54212 | M-021231-01 |
| **RIPK1** | 8737 | M-004445-02 | **SLC25A11** | 8402 | M-007470-00 | **SOD2** | 6648 | M-009784-02 |

### Table S1

| | | | | | | | | |
|---|---|---|---|---|---|---|---|---|
| **RLBP1L1** | 157807 | M-018542-01 | **SLC25A12** | 8604 | M-007471-01 | **SORD** | 6652 | M-008323-02 |
| **RNF185** | 91445 | M-007107-01 | **SLC25A13** | 10165 | M-007472-01 | **SOX10** | 6663 | M-017192-00 |
| **RNF34** | 80196 | M-007072-00 | **SLC25A14** | 9016 | M-007473-00 | **SPATA18** | 132671 | M-016038-01 |
| **RNF5** | 6048 | M-006558-02 | **SLC25A15** | 10166 | M-007474-02 | **SPATA19** | 219938 | M-017842-00 |
| **ROBO3** | 64221 | M-026504-00 | **SLC25A16** | 8034 | M-007475-00 | **SPG20** | 23111 | M-015681-02 |
| **ROM1** | 6094 | M-017459-00 | **SLC25A17** | 10478 | M-007476-01 | **SPG7** | 6687 | M-006039-02 |
| **RP2** | 6102 | M-012350-01 | **SLC25A18** | 83733 | M-007477-01 | **SPIN1** | 83985 | M-003987-03 |
| **RPE65** | 6121 | M-008199-01 | **SLC25A19** | 60386 | M-007478-02 | **SPTLC1** | 10558 | M-006673-02 |
| **RPS3** | 6188 | M-013607-01 | **SLC25A2** | 83884 | M-007479-01 | **SQRDL** | 58472 | M-008271-01 |
| **RPS6KB1** | 6198 | M-003616-03 | **SLC25A20** | 788 | M-007480-03 | **SRC** | 6714 | M-003175-03 |
| **RPS6KC1** | 26750 | M-005371-02 | **SLC25A21** | 89874 | M-007481-01 | **STAR** | 6770 | M-019369-01 |
| **RUFY2** | 55680 | M-007016-01 | **SLC25A22** | 79751 | M-007482-01 | **STARD13** | 90627 | M-010256-00 |
| **RUNX1** | 861 | M-003926-02 | **SLC25A23** | 79085 | M-007360-01 | **STARD3** | 10948 | M-017665-00 |
| **RUNX2** | 860 | M-012665-01 | **SLC25A24** | 29957 | M-007325-01 | **STARD7** | 56910 | M-017289-01 |
| **RXRA** | 6256 | M-003443-02 | **SLC25A25** | 114789 | M-007355-01 | **STAT2** | 6773 | M-012064-00 |
| **S100A8** | 6279 | M-011770-02 | **SLC25A26** | 115286 | M-007343-01 | **STAT5A** | 6776 | M-005169-02 |
| **S100A9** | 6280 | M-011384-02 | **SLC25A27** | 9481 | M-007483-01 | **STAT5B** | 6777 | M-010539-02 |
| **S100B** | 6285 | M-012259-02 | **SLC25A28** | 81894 | M-007368-01 | **STIM1** | 6786 | M-011785-00 |
| **STK32A** | 202374 | M-004634-01 | **TP73** | 7161 | M-003331-01 | **WASF1** | 8936 | M-011557-01 |
| **STOM** | 2040 | M-016971-00 | **TPM1** | 7168 | M-017837-01 | **WFS1** | 7466 | M-009740-01 |
| **STOML2** | 30968 | M-020518-00 | **TPRT** | 23590 | M-008464-01 | **WWOX** | 51741 | M-003961-03 |
| **STOX1** | 219736 | M-017070-01 | **TPT1** | 7178 | M-004559-03 | **XBP1** | 7494 | M-009552-02 |
| **STX17** | 55014 | M-020965-01 | **TRAF2** | 7186 | M-005198-00 | **YBX1** | 4904 | M-010213-03 |
| **SUCLG1** | 8802 | M-008677-01 | **TRAK1** | 22906 | M-020331-02 | **YIPF1** | 54432 | M-016906-00 |
| **SUCLG2** | 8801 | M-008918-01 | **TRAP1** | 10131 | M-010104-01 | **YME1L1** | 10730 | M-006103-01 |
| **SURF1** | 6834 | M-011786-01 | **TREM1** | 54210 | M-017974-00 | **YRDC** | 79693 | M-018139-01 |
| **SYNJ2BP** | 55333 | M-021176-01 | **TRIAP1** | 51499 | M-020809-00 | **YWHAB** | 7529 | M-008766-02 |
| **TARDBP** | 23435 | M-012394-01 | **TRPC3** | 7222 | M-006509-01 | **YWHAE** | 7531 | M-017302-02 |
| **TARP** | 445347 | M-032367-02 | **TRPM2** | 7226 | M-004193-02 | **YWHAG** | 7532 | M-008844-00 |
| **TAZ** | 6901 | M-009608-00 | **TRPM8** | 79054 | M-006517-01 | **YWHAH** | 7533 | M-010626-01 |
| **TBC1D5** | 9779 | M-020775-01 | **TRPV4** | 59341 | M-004195-00 | **YWHAQ** | 10971 | M-012329-00 |
| **tcag7.1260** | 441282 | M-032608-01 | **TSPAN6** | 7105 | M-010624-02 | **YWHAZ** | 7534 | M-003332-01 |
| **TCHP** | 84260 | M-014843-01 | **TSPO** | 706 | M-009559-03 | **ZCCHC17** | 51538 | M-016869-00 |
| **TECTA** | 7007 | M-012065-01 | **TST** | 7263 | M-010120-00 | **ZNF205** | 7755 | M-019556-01 |
| **TEGT** | 7009 | M-004118-01 | **TTC19** | 54902 | M-013361-01 | **Bax** | 581 | M-003308-03 |
| **TF** | 7018 | M-011189-00 | **TTC8** | 123016 | M-021417-01 | **OPA1** | 4976 | M-005273-00 |
| **TFAP2A** | 7020 | M-006348-03 | **TTR** | 7276 | M-012554-02 | | | |
| **TFDP1** | 7027 | M-003327-04 | **TUBA1A** | 7846 | M-013150-00 | | | |
| **TFR2** | 7036 | M-009686-02 | **TUFM** | 7284 | M-016741-00 | | | |
| **TFRC** | 7037 | M-003941-02 | **TULP1** | 7287 | M-011413-01 | | | |
| **TGM2** | 7052 | M-004971-00 | **TUSC2** | 11334 | M-006478-00 | | | |
| **THEM4** | 117145 | M-008604-01 | **TXLNA** | 200081 | M-017959-02 | | | |
| **TIAM1** | 7074 | M-003932-02 | **TXN2** | 25828 | M-017448-00 | | | |
| **TIMM10** | 26519 | M-008209-01 | **TXNRD1** | 7296 | M-008236-02 | | | |
| **TIMM13** | 26517 | M-009538-02 | **TYMS** | 7298 | M-004717-02 | | | |
| **TIMM17A** | 10440 | M-012739-01 | **U5-200KD** | 23020 | M-014161-00 | | | |
| **TIMM17B** | 10245 | M-020047-00 | **UBB** | 7314 | M-013382-01 | | | |
| **TIMM22** | 29928 | M-006473-02 | **UBIAD1** | 29914 | M-020412-01 | | | |
| **TIMM44** | 10469 | M-003864-00 | **UCP1** | 7350 | M-007636-01 | | | |
| **TIMM50** | 92609 | M-023692-01 | **UCP2** | 7351 | M-005114-00 | | | |
| **TIMM8A** | 1678 | M-010342-01 | **UCP3** | 7352 | M-007638-01 | | | |
| **TIMM8B** | 26521 | M-008491-00 | **UCRC** | 29796 | M-010899-01 | | | |
| **TIMM9** | 26520 | M-009572-00 | **ULK1** | 8408 | M-005049-00 | | | |
| **TK2** | 7084 | M-006788-03 | **UNC84A** | 23353 | M-025277-01 | | | |
| **TMC1** | 117531 | M-017206-01 | **UNQ9438** | 387990 | M-032055-00 | | | |
| **TMEFF2** | 23671 | M-010654-00 | **UQCC** | 55245 | M-017986-00 | | | |
| **TMEM102** | 284114 | M-018245-00 | **UQCR** | 10975 | M-032278-00 | | | |
| **TMEM126A** | 84233 | M-014829-00 | **UQCRB** | 7381 | M-015787-00 | | | |
| **TMEM126B** | 55863 | M-018904-00 | **UQCRC1** | 7384 | M-004748-00 | | | |
| **TMEM127** | 55654 | M-020232-01 | **UQCRC2** | 7385 | M-008334-02 | | | |
| **TMEM14A** | 28978 | M-018878-01 | **UQCRFS1** | 7386 | M-020100-01 | | | |
| **TMEM14C** | 51522 | M-020269-02 | **UQCRH** | 7388 | M-020101-02 | | | |

#### Table S1

| | | | | | |
|---|---|---|---|---|---|
| **TMEM16F** | 196527 | M-003867-01 | **UQCRQ** | 27089 | M-012517-01 |
| **TMEM173** | 340061 | M-024333-00 | **USH1C** | 10083 | M-020028-02 |
| **TMEM192** | 201931 | M-016802-01 | **USH2A** | 7399 | M-012381-00 |
| **TMEM203** | 94107 | M-015191-00 | **USMG5** | 84833 | M-014982-01 |
| **TMEM39A** | 55254 | M-020434-01 | **USP30** | 84749 | M-021294-03 |
| **TMEM39B** | 55116 | M-021133-00 | **USP8** | 9101 | M-005203-01 |
| **TMEM70** | 54968 | M-015366-01 | **UTRN** | 7402 | M-012382-01 |
| **TMIE** | 259236 | M-016265-01 | **VAMP1** | 6843 | M-012497-00 |
| **TMLHE** | 55217 | M-031958-00 | **VAPB** | 9217 | M-017795-00 |
| **TMPO** | 7112 | M-027195-02 | **VARS2** | 57176 | M-008268-01 |
| **TNFRSF10B** | 8795 | M-004448-00 | **VAT1** | 10493 | M-009726-01 |
| **TNFRSF5** | 958 | M-008101-02 | **VAV1** | 7409 | M-003935-01 |
| **TNFSF10** | 8743 | M-011524-01 | **VCL** | 7414 | M-009288-01 |
| **TOMM20** | 9804 | M-006487-00 | **VCP** | 7415 | M-008727-01 |
| **TOMM22** | 56993 | M-015878-01 | **VDAC1** | 7416 | M-019764-00 |
| **TOMM34** | 10953 | M-016439-00 | **VDAC2** | 7417 | M-019766-00 |
| **TOMM40** | 10452 | M-012732-00 | **VDAC3** | 7419 | M-020850-01 |
| **TOMM40L** | 84134 | M-016907-01 | **VEGFB** | 7423 | M-015731-01 |
| **TOMM7** | 54543 | M-013762-00 | **VHL** | 7428 | M-003936-00 |
| **TOMM70A** | 9868 | M-021243-01 | **VISA** | 57506 | M-024237-02 |
| **TP53** | 7157 | M-003329-03 | **VRK2** | 7444 | M-004684-02 |

**Table S2**

| siRNAs that increase % **Bax-positive** | siRNAs that increase % **Bax-negative** | siRNAs that increase % **Bax/Omi-positive** |
|---|---|---|
| **Plate1** | | |
| MARCH5 9.7 | EIF3A 226.0 | BLOC1S1 9.5 |
| C19orf12 15.3 | SLC25A46 198.0 | C19orf12 16.3 |
| BLOCS1 16.7 | BNIP3 196.0 | ATCAY 23.7 |
| PINK1 23.0 | DYNLL2 195.0 | GPER1 24.0 |
| MERTK 39.0 | FMR1 192.0 | **BNIP3L** 30.0 |
| ATCAY 43.3 | GLTSCR2 185.0 | MRPL12 33.0 |
| SIRT3 54.7 | RAB11FIP1 189.0 | IER3 37.0 |
| CD68 55.3 | RAP1B 183.0 | NR2C2 38.7 |
| KLHL7 58.3 | OPTN 170.0 | **RMDN3** 39.5 |
| STYXL1 58.7 | MTFP1 167.0 | ULK1 48.3 |
| RNF5 65.0 | TRAK2 155.0 | DISC1 50.3 |
| NR2C2 66.3 | | FUNDC1 59.3 |
| | | MARCH5 64.0 |
| | | LETM1 78.8 |
| | | RNF5 85.3 |
| **Plate 2** | | |
| GNB2L1 17.7 | PYCR1 137.0 | TRAK1 7.0 |
| SPATA19 15.7 | AARS 136.0 | SPATA19 15.0 |
| TMEM127 23.0 | SLC44A1 135.0 | TRIAP1 22.0 |
| ACADL 32.7 | TIMM50 133.7 | TMEM127 24 |
| PHB 37.0 | PXN 121.7 | PHB 33.25 |
| FAHD1 38.0 | SLC25A36 110.8 | SLC25A14 33.3 |
| PLSCR3 44.33 | | **ATG12** 42.75 |
| ATG12 47.75 | | |
| PRELID1 56.0 | | |
| TRIAP1 69.25 | | |
| PLA2G6 62.25 | | |

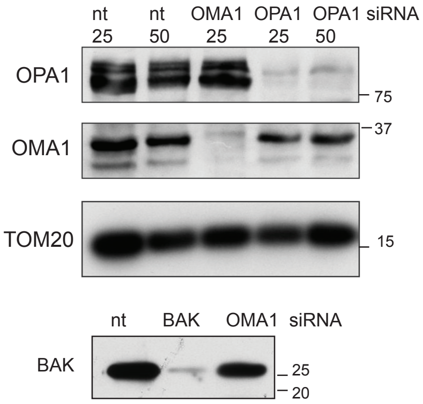
nt 25
nt 50
OMA1 25
OPA1 25
OPA1 50
siRNA
OPA1
75
OMA1
37
TOM20
15
nt
BAK
OMA1
siRNA
BAK
25
20

**Figure S1**

**A**

| sgRNA | Sequence | Exon | ICE % indels | KO score |
|---|---|---|---|---|
| **BNIP3L** | GUUCAUGGGUAGCUCCACCC | 2 | 83 | 74 |
| **RMDN3** | CUCCUGCGGGCUGUCCCAGG | 2 | 94 | 68 |
| **ATG12** | GUUUCAUACCAACUGUUCUG | 2 | 97 | 89 |

**B**

**RMDN3 CRISPR pool**

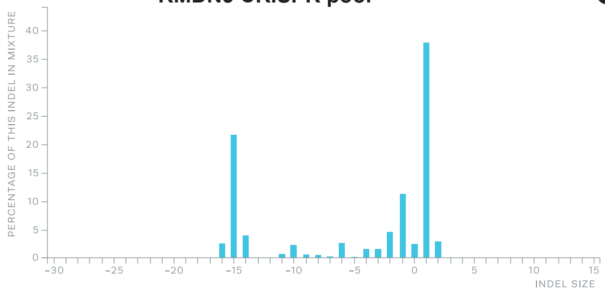


**C**

**RMDN3 CRISPR clone**

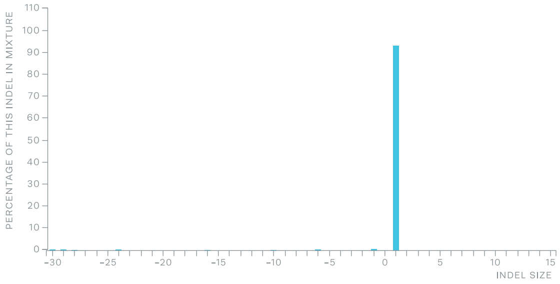


**Figure S2**
